# Changes in Protein Turn-over rates Regulate Tryptophan and Glucosinolate Biosynthesis, IAA Transport and Photosynthesis in *Arabidopsis* Growth and Defense Transitions

**DOI:** 10.1101/2023.03.31.535076

**Authors:** Mohammad Abukhalaf, Carsten Proksch, Domenika Thieme, Jörg Ziegler, Wolfgang Hoehenwarter

## Abstract

An organism continuously experiences shifts in biological states necessitating extensive rearrangement of physiology and molecular order of the cell. Here we model transitions between optimal growth conditions (homeostasis), fully induced pattern triggered immunity (PTI) and back in *Arabidopsis thaliana*, chronologically measuring changes in protein synthesis and degradation rates, transcript, protein and phytohormone abundance of 99 targets using qPCR and LC-MS parallel reaction monitoring (PRM). Temporally changing synthesis and degradation rates were primary determinants of abundance, next to changes in mRNA levels, of tryptophan, glucosinolate (GS) biosynthesis and photosynthesis associated (PAP) proteins particularly in the earlier establisher phases but also in fully induced PTI. While transcripts returned to growth levels 3 to 16 hours post elicitation, protein levels remained at fully induced PTI levels up to 16 hours into the transitory phase back to optimal growth. A notable exception were polar auxin transporters PIN3 and PIN7 levels which decreased in PTI but quickly returned to initial homeostasis levels after transition, although global auxin levels only decreased by 20%. Tryptophan, GS and JA biosynthesis proteins all increased in abundance in the wild type and the *myc234* mutant background linking induction of the tryptophan and GS biosynthesis pathways to flg22 treatment and PTI independent of MYC2 and homologs. PAPs abundance was depleted in fully induced PTI however not in the *myc234* mutant linking this active immune response to these bHLH transcription factors. FERREDOXIN-NADP(+)-OXIDOREDUCTASE (FNR1) synthesis rates decreased while its degradation rate increased in the earlier PTI establisher phase. FNR1 is the penultimate protein in the photosynthetic electron transfer chain and imparts electrons onto NADP^+^ however in its absence electrons are used for oxygen photoreduction and H_2_O_2_ production, an active defense compound. Thus FNR1 may be a molecular switch that switches photosystem activity between growth and defense under post-transcriptional control. The *myc234* mutation generally led to delayed changes in transcript and protein abundance and also abolished IAA depletion. Protein turn-over rates of a set of PAPs were affected in the mutant suggesting a possible positive role of the transcription factors in controlling post-transcriptional regulatory processes in PTI induction.

## Introduction

Plants are sessile organisms, and their life is characterized by the struggle to maintain a state of optimal growth and propagation, termed homeostasis, in a constantly changing biotic and abiotic environment. Interactions with the environment inevitably bring with them shifts to states of stress or alternative development or growth, characterized by shifting physiological dynamic equilibria.

The state transitions require effective intra- and intercellular communication. In plants, phytohormones and small peptides are among the primary molecular messengers. Plant growth is primarily regulated by auxin/IAA, gibberellic acid (GA), cytokinins and brassinosteroids. Jasmonic acid (JA) in conjunction with ethylene (ET) and salicylic acid (SA) are the two antagonistic branches that form the backbone of plant immunity, the former two to insect and necrotrophic pathogens, the latter to biotrophic pathogen attack (*1, 2*). Research in the last ten years however has made it evident, that all phytohormones contribute to both growth and developmental processes as well as biotic and abiotic interactions (*3, 4*).

Plant growth is essentially composed of stem cell division and differentiation in the apical meristems followed by cell elongation (*5*). It is orchestrated by a complex genetic program integrating diverse endogenous and environmental cues, including light, sugar and temperature as well as phytohormone signaling entailing differential expression of thousands of genes. Auxin influences all aspects of plant development and growth, especially cell division and elongation (*6*). Auxin/IAA is primarily synthesized from tryptophan by way of four pathways in *Arabidopsis thaliana*, utilizing the tryptophan metabolic products indole-3-acetaldoxime (IAOx), indole-3-acetamide (IAM), indole-3-pyruvic acid (IPyA), and tryptamine (TAM) as substrates, respectively (*7*). It is synthesized in the shoot, root and young leaf apical meristems and then transported from these regions of highest concentration throughout the plant by two transport systems, non-directional phloem stream and directional polar cell-to-cell transport (PAT) (*7*). PAT is facilitated by membrane localized auxin efflux carriers of the PIN-FORMED family (PIN). PIN1 transports auxin in a basipetal direction from shoot to root whereas it is transported acropetally by PIN 2 (*8*). The other canonical PINs PIN3, PIN4 and PIN7 transport the phytohormone away from tissue apices forming a type of “reflux loop” (*9*), making auxin patterning and establishment of local concentration gradients essential regulators of growth and developmental processes in all plant organs.

Plants have a decentralized immune system that induces pathogen associated molecular pattern (PAMP)/pattern recognition receptor (PRR) triggered immunity (PTI) upon perception of PAMPs by cell surface plasma membrane spanning receptors (PRRs) (*10*). This is exemplified by the PRR receptor kinase (RK) FLAGELLIN SENSITIVE 2 (FLS2) binding the 22 amino acid N-terminal epitope of bacterial flagellin (flg22) (*11*). Subsequent assembly of the mature, active receptor complex by recruitment of accessory receptor-like proteins (RLPs; BAK1 and others) (*12, 13*) and scaffolds (RK FERONIA family and others) (*14*) initiates phosphorylation mediated signaling. It is propagated by receptor-like cytoplasmic kinases (RLCKs) such as BOTRYTIS-INDUCED KINASE 1 (BIK1) and PBS1-like kinases (*12*) that phosphorylate calcium channel proteins to promote Ca^2+^ influx (*15*), calcium-dependent protein kinases such as CPK5 and NADPH OXIDASE RESPIRATORY BURST OXIDASE HOMOLOG (RBOHD). RBOHD phosphorylation by CPK5 (*16*) and Ca2+ binding allows the protein to induce the ROS burst. Upstream components of MAP kinase (MAPK) cascades are also phosphorylated and activated by RLCKs (*17*), which in turn target 100s of downstream substrates (*18*). CPKs (*19*) and MPKs converge on transcription factors most notably SARD1 that regulate expression of *ics1* and SA biosynthesis. All of these partially interconnected signaling relays ultimately lead to differential expression of thousands of genes (*20*) and extensive proteome remodeling (*21*). Among others, PR proteins with anti-microbial activity (*22*) increase in abundance. Proteins in secondary metabolites/defense compounds synthesis pathways such as glucosinolates (GS) also increase in their abundance. Indolic glucosinolates (IGs) are synthesized from tryptophan (*23*) and upon cleavage by thioglucosidases called myrosinases give rise to a wide variety of toxic compounds such as isothiocyanates, thiocyanates and nitriles (*24, 25*).

Salicylic acid (SA) and SA signaling are central and integrative components of both PTI and nucleotide like receptor (NLR)/effector triggered immunity (ETI) (*26*). Suppression of JA levels by SA in biotrophic pathogen resistance is well known (*1*), however a role of JA in PTI (*27–30*) and complementary activity of both hormones in fine tuning immunity to divergent pathogen lifestyles (*2*) have been documented, as well as numerous JA biosynthetic and signaling genes responding to flg22 (*20*). The first step in JA biosynthesis is the release of α-linolenic acid (α-LeA) from chloroplast membranes. After several oxidative enzymatic reactions, the JA intermediate 12-oxo-phytodienoic acid (OPDA) is exported from chloroplasts and further reduced in peroxisomes to (+) 7-iso-JA. JA is then exported into the cytoplasm. The bioactive compound JA-Isoleucine (JA-Ile) produced by conjugation of isoleucine to JA by the jasmonate-isoleucine synthetase JAR1 is perceived by the SCF^COI1^–JAZ co-receptor complex. This leads to the degradation of transcriptional repressor JAZ proteins thus liberating expression of genes under the control of the JA signaling pathway. These include the basic helix-loop-helix (bHLH) transcription factor MYC2 and its homologs MYC3, MYC4 and MYC5, transcriptional activators that extensively orchestrate the genetic JA response program (*31*). The JA signaling cascade and MYC2 activity are preeminent in the context of wounding and defense against biting insects but studies in the past years have elucidated a function in the response to flg22 and immunity to biotrophic pathogens (*3, 27, 28, 32*). MYC2 is a master regulator of gene expression and a central hub in the larger network that integrates phytohormone cross-talk and signaling (*33*). It has been shown to bind to more than 6,000 genes and more than 60% of all genes responding to exogenous JA application (*34*). Furthermore, nearly half of 1,717 known or predicted *Arabidopsis thaliana* TF genes were direct MYC2/3 targets and/or JA responsive. Also, the expression of many JA pathway genes including JA synthesis itself is under the control of MYC2 and MYC2 abundance and activity are extensively auto-regulated (*35–38*), evidence of positive and negative feedback control (*39*).

Photosynthesis is the process of atmospheric carbon assimilation to produce higher energy organic biomolecules. It generates a strong reducing potential by way of transfer of electrons produced by oxidation of water and oxygen evolution across chloroplast thylakoid membrane integral photosystems II and I (PSII and PSI). The molecular machinery is composed of a plethora of proteins and other biomolecules and the abundance of these proteins is diminished upon induction of PTI by flg22 (*21, 40*). This is an active defense response, the meaning of which however is not fully understood.

Cellular proteins undergo constant turn-over (*41*). The protein pool is replenished by new synthesis and depleted by degradation of old proteins. The rate of synthesis is considered to follow zero order kinetics meaning it is independent of the amount of protein present. The degradation rate is modelled as first order kinetics meaning it depends on the amount of protein present in the pool over time. In physiological steady states such as homeostasis, the two are in equilibrium and any changes in protein abundance are only due to growth. Shifts between steady states such as between growth and immunity lead to changes in protein abundance due to proteome remodeling and perturb this equilibrium, resulting in potentially altered synthesis and degradation rates. Research in the last years have produced evidence that translational mechanisms play substantial roles in regulating plant immunity (*42–44*). Changes in synthesis and degradation rates of proteins may be one such control mechanism.

Medium to large scale measurements of synthesis and degradation rates are possible combining a partial stable heavy isotope protein labeling strategy and mass spectrometry based proteomics (*45*). Under laboratory conditions the organisms under study are switched to a stable heavy isotope containing diet as anabolic precursors at a given stage of development or growth. Incorporated into proteins, this leads to a shift in the peptide ion isotope envelope to greater mass to charge ratios (m/z) in liquid chromatography mass spectrometry (LC-MS) measurements of tryptic peptides over time. Increased signal intensity of higher order isotope peaks is indicative of new protein synthesis due to incorporation of the heavy isotope into peptide primary structure whereas quenching of the naturally predominant monoisotopic peak is the result of protein degradation. Labeling proteins with the heavy nitrogen isotope ^15^N has proven to be the method of choice in plants (*46*) and has been demonstrated successfully in *Arabidopsis* (*47*), barley (*48*) and *Medicago truncatula* (*49*).

We developed a chronological quantitative proteomics model of PTI comprising the tryptophan, GS, JA biosynthesis pathways, members of the auxin biosynthesis pathway and phytohormone conjugation and metabolism and auxin and JA signaling previously (*21*). Additional components were photosynthesis associated proteins (PAPs) and proteins playing roles in primary metabolism. Here we expanded our research interest to chronological changes in the abundance of these 99 proteins and the underlying regulatory aspects of the model in the shifts between the steady states of homeostasis to fully induced PTI and back. To this end, *Arabidopsis* seedlings were grown in liquid culture for ten days and then supplemented with flg22 (1 µM in medium). Plants were sampled at the time of induction, 1, 3 and 16 hours post induction, followed by transfer of plants back to flg22 free medium and sampling 1, 3 and 16 hours post transfer (later referred to as 17, 19 and 32 hrs). Proteins, mRNA and metabolites were extracted from plant tissue and measured using parallel reaction monitoring (PRM) targeted proteomics, qPCR and multiple reaction monitoring (MRM), respectively. Proteins synthesis and degradation rates (K_s_ and K_d_) were also measured to further elucidate post-transcriptional control mechanisms. The *myc234* knockout mutant was added to further explore the role of JA and JA signaling in PTI.

## Materials and Methods

Plant cultivation, growth, treatment, sample preparation, targeted PRM LC-MS measurements, peptide and protein identification, PRM based area under the curve (AUC) peptide and protein quantification, mRNA isolation, cDNA synthesis and qPCR (Supplementary table Primers) and phytohormone extraction, sample preparation and MRM LC-MS phytohormone measurements were performed as described previously in the open access paper by Bassal and co-workers (*21*).

### Measurement of flg22

Media samples at time points 0, 1, 16, 17 and 19 hrs. were filtered with Amicon® Ultra 30K filters. Filters were washed overnight in 5% Tween-20 and then three times with ddH_2_O for 30 minutes each. 300 µL of the media sample were added to the filter and then centrifuged at 16,100 g for 10 min at RT. The filtrates were kept and dried in a vacuum concentrator and the resulting dried samples were desalted on STAGE-Tips C18. The final dried peptide samples were dissolved in 136 µL 5% ACN, 0.1% TFA to reach a final concentration of 2.2 µM flg22 for LC-MS analysis.

Dried peptides were dissolved in 5% acetonitrile, 0.1% trifluoroacetic acid, and injected into an EASY-nLC 1000 liquid chromatography system. Peptides were separated using liquid chromatography C18 reverse phase chemistry employing a 60 min gradient increasing from 5% to 40% acetonitrile in 0.1% FA, and a flow rate of 250 nL/min. Eluted peptides were electrosprayed on-line into a Q Exactive^TM^ Plus mass spectrometer. The spray voltage was 1.9 kV, the capillary temperature 275°C and the Z-Lens voltage 240 V. A full MS survey scan (DDA top10) was carried out with chromatographic peak width set to 15 s, resolution 70,000, automatic gain control (AGC) 3E+06 and a max injection time (IT) of 100 ms. MS/MS scans were acquired with resolution 17,500, AGC 5E+04, IT 50 ms, loop count 10, isolation width 1.6 m/z, isolation offset 0.0 and a normalized collision energy 28.

Flg22 peptides were identified using the Mascot software v 2.5.0 (Matrix Science) linked to Proteome DiscovererTM v2.1. The enzyme was set to none. A precursor ion mass error of 5 ppm and a fragment ion mass error of 0.02 Da were tolerated in searches of the flg22 sequence as a database. Oxidation of methionine (M) was tolerated as a variable modification. A PSM, peptide and protein level false discovery rate (FDR) was calculated for all identified spectra and peptides and proteins based on the target-decoy database model. The significance threshold α was set to 0.01 to accept PSM, peptide and protein identifications.

### Measurement of protein degradation and synthesis rates

*Arabidopsis thaliana* Col-0 and *myc234* triple knockout mutant seedlings were grown in liquid culture in the following media.

|  |  |
| --- | --- |
| **25% MS medium**<br>**1L** | 0.748 mM CaCl<sub>2</sub>, 0.313 mM KH<sub>2</sub>PO<sub>4</sub>, 0.375 mM MgSO<sub>4</sub>, 4.7 mM KNO<sub>3</sub>(475 mg), 5.15 mM NH<sub>4</sub>NO<sub>3</sub>(413 mg), 2.35 mM MES buffer, 0.25x Basal Salt Micronutrients solution, 0.25x Vitamin solution (pH 5.7) |
| **25% MS medium <sup>15</sup>N**<br>**1L** | 0.748 mM CaCl<sub>2</sub>, 0.313 mM KH<sub>2</sub>PO<sub>4</sub>, 0.375 mM MgSO<sub>4</sub>, 4.7 mM K<sup>15</sup>NO<sub>3</sub>(480 mg), 5.15 mM <sup>15</sup>NH<sub>4</sub><sup>15</sup>NO<sub>3</sub>(423 mg), 2.35 mM MES buffer, 0.25x Basal Salt Micronutrients solution, 0.25x Vitamin solution (pH 5.7) |

250 *Arabidopsis thaliana* seeds were grown in 50 ml 25% MS medium on an orbital shaker 45 rpm at 22°C under long day conditions (16 hrs light, 8 hrs dark) for ten days. Cultures were harvested in triplicate representing the start of the experiment t=0. Remaining cultures were transferred to 25% MS medium containing ^15^N as the sole nitrogen source and grown further under the same conditions. Cultures were sampled in triplicate 8, 9, 10, 12, 24, 36, 48, 72 and 96 hrs after media exchange to measure protein synthesis and degradation rates under optimal growth conditions.

To measure protein synthesis and degradation rates under PTI conditions, flg22 was added to a concentration of 1 µM in medium, 8 hrs after exchange to ^15^N medium. Cultures were sampled at this time point in triplicate and further sampled 2, 4, 6, 8, 16, 24, 32, 48, 72 and 96 hrs after flg22 injection (corresponding to 10, 12, 14, 16, 24, 32, 56, 80 and 104 hrs after media exchange). Harvested tissue was weighed and frozen at −80°C.

Proteins were extracted and samples prepared for MS analysis as described in (*21*). Ion peak isotope envelopes of proteotypic target peptides of the 99 cognate target proteins were measured with the PRM strategy also described in (*21*). Raw data were converted to .MzML format using msconvert (*50*). Protover, a python-based code program was used to extract the relative isotopic abundance (RIA) of ^15^N in peptide ion isotope envelopes over all sampling time points (*51*) (equation 1).

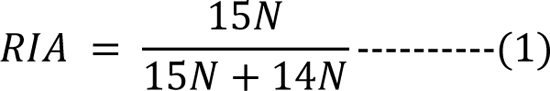

where 15N represent the heavy and 14N the naturally abundantly occurring isotope population

The list of proteotypic target peptides with their average retention times gained from the PRM experiments was used as an input for the program. A retention time window of 5 min was set as filter and the “RIA increasing” filter was disabled. The program output which consisted of target peptides and their RIA in all measurements of all replicates at all sampling time points were further analyzed using Excel. Only peptides with RIA values at 5 or more time points in at least one replicate were retained. Then median RIA values of cognate proteotypic peptides were used to infer RIA values for respective proteins at all sampling time points in each replicate.

Under optimal growth conditions the apparent degradation rate (K_loss_) was calculated for each protein from the slope of the logarithmic function -ln(1-RIA) plotted over time as in equation 2.

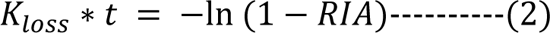

where t is time in days.

Then proteins with relative standard errors (RSTDE) of K_loss_ values of the three replicates greater than 40%, median R^2^ values below 0.5 and proteins with negative K_loss_ values were discarded. The actual degradation rate constant (K_d_) for each protein was then calculated using equation 3.

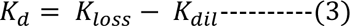

where K_dil_ is the doubling constant.

K_dil_ was calculated by fitting an exponential model to the weight increase of seedlings over experimental sampling time points as a proxy for the increase of total protein amount over time as in equation 4.

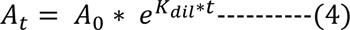

where A_t_ is the total amount of proteins at time (t) and A_0_ is the proteins amount at t=0. To calculate the synthesis rate constant (K_s_/A), equation 5 (*47*) was used.

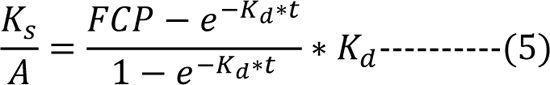

where FCP is Fold Change of Protein Abundance.

By reorganizing equation 4, FCP can be calculated as follows:

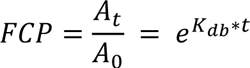

By substitution in equation 5 (K_s_/A) can be calculated as in equation 6.

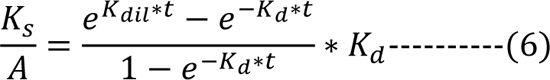

A mean time of 2 days was used to calculate K_s_/A for all filtered proteins (later as K_s_).

Protein K_s_ and K_d_ values were calculated separately for the transitionary phase between optimal growth and fully induced PTI at 16 hrs post flg22 exposure and fully induced PTI itself, 16 hrs and beyond. For the early establisher phase only time points 0, 2, 4, 6, 8 and 16 hrs after flg22 treatment were used. To calculate the K_d_, the slope of the logarithmic function in equation 7 was used:

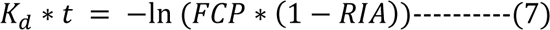

However, here FCP was estimated at the respective time points by fitting a polynomial second order model to the PRM data protein abundance measurements using median FCP at time points 0, 1, 3 and 16 hrs. Only proteins with FCPs of 1.5 or more **or** 0.75 or less at any of the time points 1, 3 or 16 hrs were considered. The same estimated PRM data FCPs were used for all biological replicates. Median peptide RIA values were used to infer cognate protein RIA values for K_d_ calculations and proteins with measured RIA values at 4 time points or more in at least one biological replicate were considered. Proteins with relative standard errors (RSTDE) of K_d_ values of the three replicates (TSA1 and AAO1 had only two replicates) equal to or less than 40% and median R^2^ values of 0.5 or more were considered. For calculation of K_d_ values in fully induced PTI equations 2 to 4 were used as above using time points 0,8,16,24,32,48,72 and 96 hrs after flg22 treatment in the K_loss_ and K_dil_ calculations. K_s_ values in both phases were calculated using equations 5 and 6 as above.

## Results

### Targeted multi-omics experiment of shifts between growth and immunity

231 peptides derived from the 99 model proteins were quantified in a single run using retention time scheduled PRM (*52*). These peptides were proteotypic for the 99 proteins, the abundance of each protein being monitored with at least 1 proteotypic peptide (Supplementary table 1). The transcript levels of 34 proteins from the same set were measured with an extra 4 proteins which were not amenable to MS measurements included: CYTOCHROME P450 FAMILY 94 SUBFAMILY B POLYPEPTIDE 1 and 3 (CYP94B1 and CYP94B3), CYTOCHROME P450 FAMILY 94 SUBFAMILY C POLYPEPTIDE 1 (CYP94C1) and ISOCHORISMATE SYNTHASE 1 (ICS1) (Supplementary table 2). Moreover, we quantified the phytohormones abscisic acid (ABA), Indole-3-acetic acid (IAA), JA, JA-Ile, 12-hydroxy-JA (12-OH-JA), (+)-12-oxo-phytodienoic acid (OPDA) and SA which have important roles in growth and/or defense (Supplementary table 3).

### Experimental design validation

We wanted to control that no flg22 is transferred with the seedlings meaning the recovery phase back to homeostasis is clear of elicitor. To this end we quantified flg22 in growth media at sampling time points 0, 1, 16, 17, and 19 hrs (Supplementary Fig 1) (Supplementary table 4). As expected flg22 was not detected after transfer of the seedlings, however, we were surprised to find that one hour after introduction to the medium full length flg22 accounted for only 9% of all identified flg22 derived peptides (breakdown products) and after 16 hours the full length peptide was depleted completely and only three products with 10, 11 and 12 amino acids remained. Nonetheless, these flg22 breakdown products contain the core epitope recognized by FLS2 and thus presumably were still active, though less efficiently (*53*).

**Figure 1:**
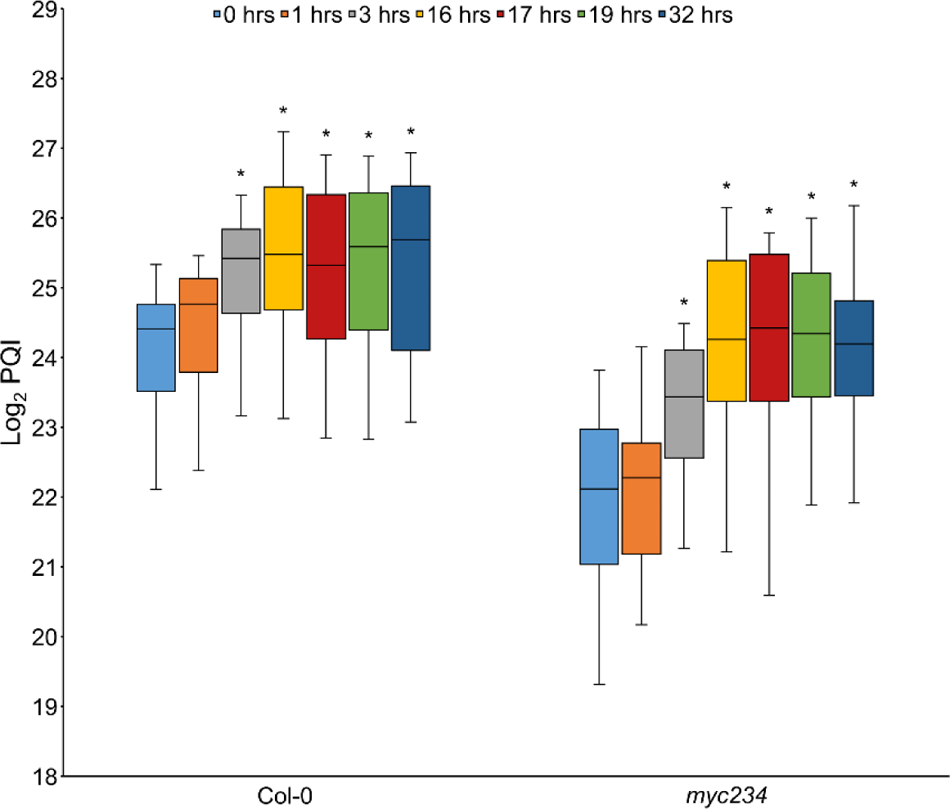
Plotted LOG_2_ protein quantification indexes (PQI) were median area under the curve (AUC) values of targeted proteotypic peptides for each cognate protein. PQI values for tryptophan,GS and other secondary metabolites biosynthesis proteins are plotted (n=18) for the different experimental sampling points. Box represents 2^nd^ and 3^rd^ quantiles, whiskers 1^st^ and 4^th^ quantiles respectively, line the median. Stars indicate statistically significant differences in respect to the 0 hrs time point as determined by a two-sided student’s t-test with α=0.05.

To ensure that the physical transfer did not elicit changes in protein abundance, we performed an experiment in which plants were mock treated with water and samples were taken at 0, 16, 17 and 19 hrs in quadruplicate. We quantified our target proteins and analyzed the results using hierarchal clustering (Supplementary Fig 2A). Both row and column dendrograms showed random clustering. Proteins produced two clusters, the major cluster containing 92 of the 99 target proteins (Supplementary Fig 2B). Additionally, log_2_ protein fold changes of sampling time points in respect to time point 0 were tested for significance using a one sample T-test (FDR multiples testing corrected α=0.01) with the null hypothesis H_0_ being no changes in protein abundance across all time points (Supplementary table 5). Only one protein (PAL1) showed a significant increase in abundance at the 16 hrs sampling point all together indicating that neither physical transfer of plants or other experimental factors including the time of day of sampling or amount of light had affected protein abundance.

**Figure 2:**
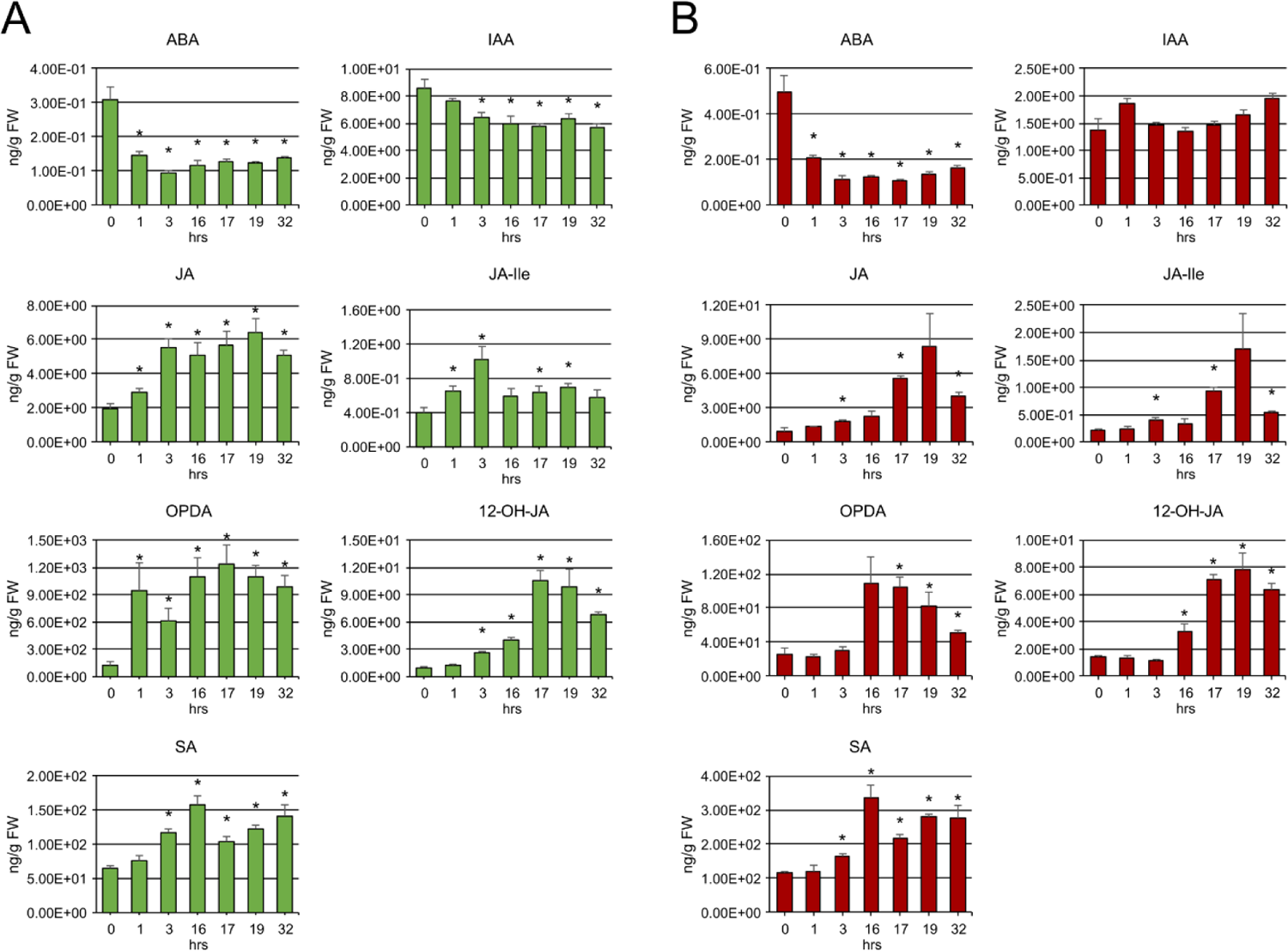
SRM based quantification of phytohormone levels throughout experimental sampling time points. Error bars denote standard error (SE). Stars indicate statistically significant differences in respect to the 0 hrs time point as determined by a two-sided student’s t-test with α=0.05 and n=4 or 3 (Col-0 or *myc234*). A. Col-0. B. *myc234* background.

### Proteome is remodeled late and remains stable into PTI recovery phase back to growth

The abundance of the 99 target proteins was measured by PRM at the indicated sampling points (Supplementary table 6). Protein levels generally increased at 1 and 3 hrs and reached maximum values at 16 hrs post elicitation which we considered to be fully induced PTI. This held particularly true for the proteins in the tryptophan biosynthesis pathway, PHOSPHORIBOSYLANTHRANILATE TRANSFERASE 1 (PAT1), INDOLE-3-GLYCEROL PHOSPHATE SYNTHASE (IGPS), TRYPTOPHAN SYNTHASE ALPHA CHAIN (TSA1) and TRYPTOPHAN SYNTHASE BETA-SUBUNIT 1 (TSB1) increasing to more than double their amounts measured in steady state growth. The same trend was observed for proteins of the GS, phenylpropanoid and 4-OH-ICN biosynthesis pathways (Fig 1) as well as several proteins involved in auxin biosynthesis such as the aldehyde oxidases. Tryptophan channels into both IG and auxin synthesis and these pathways are known to be induced upon exposure to flg22 (*23*). Interestingly, protein abundance remained elevated after transfer back to flg22 free medium until the last sampling time point at 32 hrs. This was contrary to our expectations of a return to initial levels as the plants shifted back to homeostasis from fully induced immunity.

Similar behavior was observed for target proteins that conjugate or metabolize JA and/or auxin. We reported this previously for JASMONATE-INDUCED OXYGENASE 2 (JOX2) and IAA-ALA RESISTANT3 (IAR3) in the context of a mechanistic model for control of JA and JA-Ile levels in response to flg22 (*21*). Concurrently we also measured a significant increase in 12-OH JA over the course of the shift from growth to PTI at 16 hrs post elicitation which was further amplified during the return to growth (Fig 2A, Supplementary table 3). JA and JA-Ile levels remained at nearly basal levels, increasing slightly but significantly at 1 and 3 hrs post exposure and then decreasing 16 hrs after exposure. SA levels increased significantly 3 hrs after flg22 perception and remained elevated throughout the recovery phase and the return to homeostasis. The opposite was true for ABA, whose abundance decreased significantly already after 1 hour of exposure to the PAMP and remained at approximately the same level for the rest of the sampling points of the experiment.

### Transcripts respond early to PAMP exposure and the discrepancy between transcript and protein levels

We measured the cognate transcript levels of 34 of the targeted proteins (Supplementary table 7). These showed a positive linear correlation (R^2^) of 0.36 to protein abundance in homeostasis. This correlation became pronouncedly negative after 1 hr of exposure to flg22 and deteriorated almost completely after 3 and 16 hrs. It remained negative over all PTI time points however the descent of the slope of the linear model flattened over time (linear models are shown in Supplementary table 7). This indicates transcription of the cognate genes occurs early in the steady state shift and that there is a considerable lag time before translation leads to ramping up of protein abundance. Indeed, protein levels reached a maximum 16 hours after induction of immunity. Transcript levels reached initial levels after 16 hrs of transfer to flg22 free medium.

To investigate this further we calculated the protein to transcript ratios (the amount of protein per transcript) as a proxy for synthesis of the 34 proteins as applied by Kuster and co-workers (*54*) (Supplementary table 7). The change in ratios over the experimental sampling time points was analyzed by hierarchical cluster analysis (HCL) (Fig 3A, Supplementary table 8). Two main clusters were observed, one (termed cluster A) with 16 proteins in which the ratios decreased to a minimum 1 and 3 hrs after flg22 exposure then increased well above values in unchallenged plants at the 16 hrs sampling point where PTI is fully induced. This again reflects initial elevation of transcript levels followed by a substantial lag of several hours until proteins accumulate. Ratios remained at these elevated levels throughout the recovery phase and indeed increased further at the 32 hrs sampling point where we expected return of plants to homeostasis. This shows that transcripts of the flg22 responsive proteins are degraded but that the proteins themselves remain stable even 16 hrs into the steady state shift from PTI and the return to optimal growth conditions.

**Figure 3:**
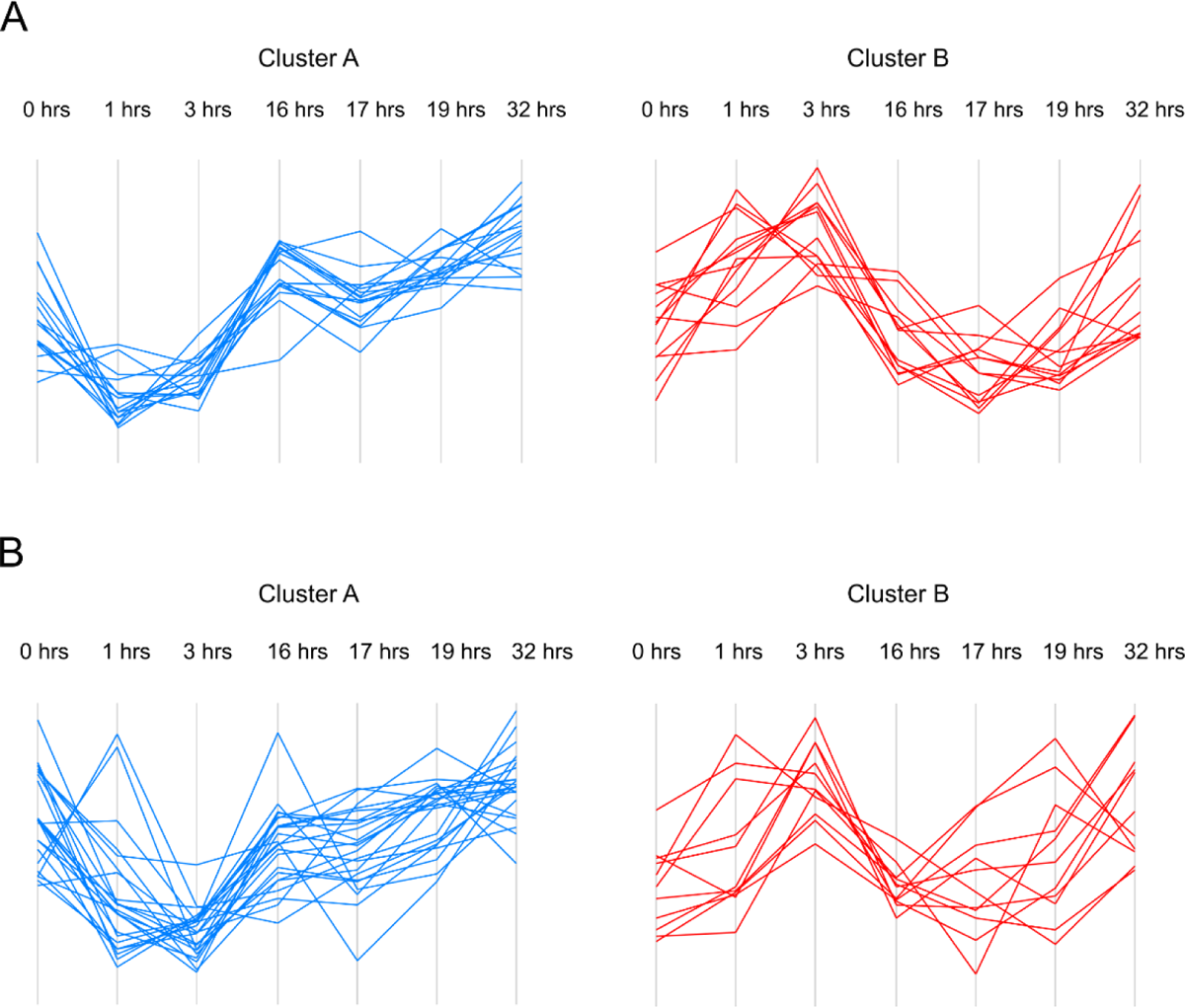
Protein (PRM based quantification) to transcript (qPCR-based quantification) abundance ratios throughout experimental sampling time points. A. Col-0. Cluster A (n=16); Cluster B (n=13). B. *myc234* background. Cluster A (n=23); Cluster B (n=11).

The second cluster (termed cluster B) contained 13 proteins whose protein to transcript ratios increased 1 and 3 hrs after challenge with the PAMP followed by a decrease to a minimum at 16 hrs post flg22 treatment and 1 hr after transfer to flg22 free medium with a subsequent return to approximately initial levels over the course of the steady state shift back to homeostasis (Fig 3A, Supplementary table 8). These proteins are hallmarked by an early decrease in the levels of their cognate transcripts followed by reduction in protein abundance several hours later. As opposed to the proteins in cluster A, the abundance of these 13 proteins increased in later stages of growth after return to the flg22 free medium, so they responded to the transition from PTI back to optimal growth in the sampled 16 hrs time frame.

These results showed a complex relationship between transcript and protein abundance presumably also governed by post-transcriptional mechanisms primarily in adaptive shifts between steady states. They also suggest that proteins accumulating under conditions of pathogen resistance especially those involved in defense compound synthesis remain abundant in the cell and that degradation and clearance of these proteins even upon return to homeostasis may not be expedient or necessary.

### Polar auxin transport may play an important role in shifts between growth and immunity

Several proteins playing roles in auxin transport and auxin and JA signaling belonged to cluster B and decreased in their abundance upon exposure to flg22. Their abundance reached a minimum at 16 hrs post induction of PTI or 1 and 3 hrs post transfer to flg22 free medium. A return to basal growth levels could however generally be observed 16 hrs after transfer to fresh medium (32 hrs time point) as opposed to the proteins whose levels were elevated in the course of PTI and remained elevated in cluster A. The auxin efflux carriers PIN3 and PIN7, which are primarily responsible for polar cell-to-cell auxin transport especially showed this trend exhibiting the greatest fold changes in abundance in this set of proteins (log_2_ fold change −1.06 and −1.45 16 hrs after flg22 treatment and max log_2_ fold change −1.28 and −1.88 1 hour after medium switch, respectively). As also reported previously (*21*) global auxin levels decreased by around 20% although significantly at α=0.05, 3 and 16 hrs post PTI induction and at the 17, 19 and 32 hrs sampling points into the recovery phase (Fig 2A, Supplementary table 3). Together this suggests auxin transport and localization may play a preeminent role in shaping PTI, in addition to reduction of the total amount of auxin/IAA itself.

### Downregulation of photosynthesis in PTI and FNR1 as a modulator between growth and defense

The abundance of 23 photosynthesis associated proteins (PAPs) involved including photosystem I and II (PSI and PSII) components that are primary orchestrators of the light reaction, Calvin cycle proteins and proteins playing roles in photorespiration were also measured (Supplementary table 6). Twenty of these 23 were significantly less abundant 16 hrs after exposure to flg22 in the wild type (Figure 4A). Protein levels remained low throughout the return to homeostasis up until the 32 hrs sampling point. Inhibition of photosynthesis has been reported previously as an active part of the plant immune response to pathogens (*55*).

**Figure 4:**
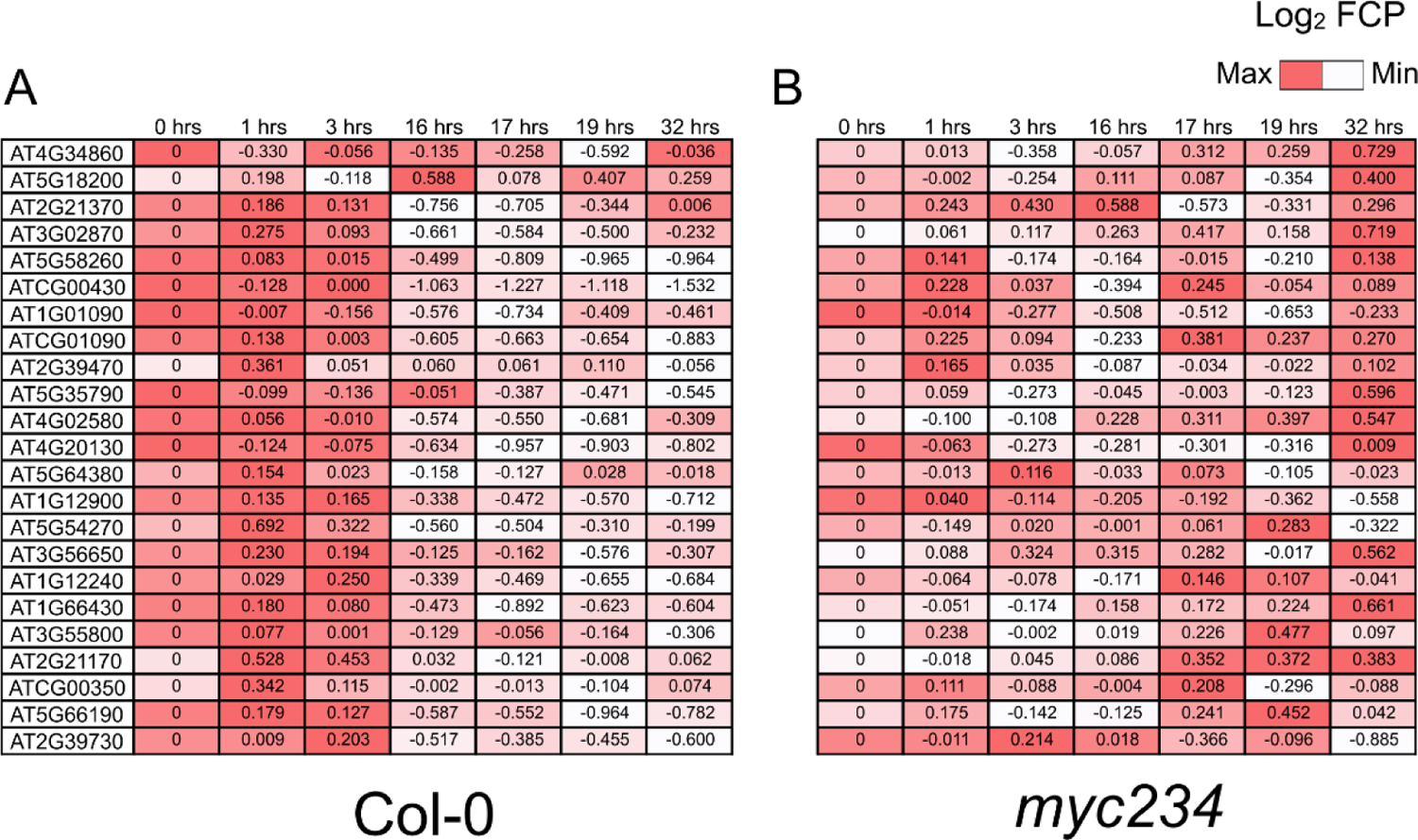
LOG_2_ fold changes in protein abundance (FCP) (PRM based quantification) in respect to the 0 hrs time point of 23 proteins involved in photosynthesis. A Col-0. B *myc234*.

Among these proteins was FERREDOXIN-NADP(+)-OXIDOREDUCTASE (FNR1). This protein is associated with PSI and catalyzes the final step in the linear electron transfer chain reducing NADP^+^ to NADPH, which goes into the Calvin cycle to produce Glyceraldehyde 3-phosphate (GAP) from atmospheric CO_2_. Notably, if photosynthetic linear electron transfer is disrupted and electrons are not transferred as readily onto NADP^+^, they are instead used to produce chloroplastic H_2_O_2_ (ROS) an integral part of plant defense (*55*). FNR1 transcripts were diminished to nearly 0.5 fold 3 hrs after PAMP exposure and remained at approximately this level throughout the remaining sampling time points (Supplementary table 2). Protein levels decreased to −0.6 log_2_ fold change 16 hrs after exposure and remained at this level or slightly lower throughout the rest of the experiment (Supplementary table 6) suggesting that the depletion of FNR1 may switch photosystem function from growth to defense.

### Delayed PTI onset, reduced auxin levels and attenuated depletion of photosynthesis proteins in *myc234* mutant

The same multi-omics experiment as in the wild type were performed in the *myc234* triple knockout mutant (*56*). The abundance of the target proteins that were constituents of the tryptophan, GS, 4-OH-ICN and phenylpropanoid biosynthesis pathways were diminished compared to the wild type (Fig 1, Supplementary Table 6). Accumulation of these proteins in the mutant was also delayed becoming pronounced at the 3 and 16 hrs time points whereas it was already higher 1 hr after exposure to the PAMP in Col-0 (Fig 1). Many of the genes in these pathways are targets of MYC2 and MYC3 (*34*) and it has been reported that their expression in the triple mutant is reduced (*57*). The abundance of proteins involved in auxin/JA conjugation and JA synthesis increased later in the *myc234* than in the wild type background, again 16 hrs as opposed to 1 and 3 hrs following exposure to flg22, respectively. On the other hand, proteins with a function in auxin and JA signaling decreased earlier in their abundance reaching a minimum at the 3 hrs as opposed to 16 and 17 hrs time points.

Concurrently, JA, JA-Ile and OPDA levels were much reduced in the triple mutant and increased more slowly if at all. 12-OH-JA was elevated in the mutant but accumulated much slower beginning at the 16 hrs time point (Fig 2B). JA synthesis, conjugation and signaling are auto-regulated by positive and negative feedback loops dependent on MYC2 (*35–38*). Auxin/IAA was less abundant in the *myc234* mutant background and levels did not change significantly throughout the experiment suggesting the moderate depletion in the wild type upon PTI elicitation is dependent on MYC2 and homologs. The amount of SA in the mutant was substantially greater, significantly accumulating to higher levels in the course of the experiment.

The linear correlation between transcript and protein levels of the 34 target proteins for which cognate transcripts were quantified was 0.41, similar to the wild type in homeostasis. However, it remained positive throughout the shift to steady state PTI albeit with a smaller slope at the 1 hrs and 3 hrs sampling points indicating a slower increase in transcript levels and later accumulation of protein at the 16 hrs sampling point when we considered immunity to be fully established (Supplementary table 7). Transcript and protein levels remained elevated at 17, 19 and 32 hrs time points as the plants shifted back to homeostasis. Protein to transcript ratios clustered similarly to the wild type although individual protein values were further removed from cluster centers (Fig 3B, Supplementary table 8). Cluster A that showed an early decrease in protein to transcript ratios contained 23 proteins in the mutant background as opposed to 16 in the wild type, cluster B that showed an early increase in ratios contained 11 as opposed to 13 proteins. Cluster members were generally conserved between wild type and mutant; 87.5% of cluster A protein to transcript ratios in the wild type were also cluster A members in the mutant, 69% of cluster B members related in this way, including the PIN3 and PIN7 auxin transporters. This indicates the same regulatory mechanisms generally apply to the expression of these genes in both genetic backgrounds. Notable exceptions that showed discordant cluster membership were AAO1, JAR1, COI1 and TPR1, proteins involved in auxin synthesis, auxin and JA conjugation and JA perception and signaling.

Interestingly, downregulation of the abundance of the proteins related to photosynthesis was nearly abolished in the *myc234* mutant background (Figure 4B), implying a role of these transcription factors in impingement of photosynthetic activity in the response to PAMP treatment. It is known that transcript levels of photosynthesis and growth related genes as well as leaf growth are repressed upon coronatine treatment, an analog of JA (*58*). Together our experiments in the triple mutant suggest a positive role of MYC2 in the establishment of PTI especially in regulation of the function of the photosystem.

### Protein turn-over regulates shifts between growth and immunity

We were interested in possible changes in protein K_s_ (synthesis) and K_d_ (degradation) rates affecting changes in protein abundance in flg22 induced PTI. We employed a stable heavy nitrogen isotope based partial protein labeling strategy (*48*), incorporating an 8-hour lag time after shifting cultures to the heavy isotope containing medium to ensure incorporation of the ^15^N label into primary structure for LC-MS measurement. This was designated as t_0_ and the start of the experiment. Culture fresh weights were recorded at the time of labeling, t_0_ and 1, 2, 4, 16, 28, 40, 64 and 88 hours into the experiment and target protein peptide ion isotope envelopes monitored at these time points. Heavy and natural isotope peak intensity values were extracted using the Protover software (*51*) (Supplementary table 9) while the dilution factor, which corrects for changes in protein abundance over time resulting from cell growth (K_dil_) was determined by model fitting to culture fresh weights (Supplemental Fig 3A, equation 4). K_d_ and K_s_ values under optimal growth/homeostasis conditions (C1) were thus calculated for 68 model proteins using equations 2 and 3 and 5 and 6 respectively, after quality control filtering (Supplementary table 10). Degradation rates were classified as slow (<0.055 day^-1^), intermediate (0.055-0.22 day^-1^) and fast (>0.22 day^-1^) as described before (*47*). These were 33, including 11 of the 23 monitored photosynthesis related proteins, 27 and 8 proteins, respectively (Supplementary table 10).

In a second experiment, flg22 was added to the culture medium (1µM concentration in medium) at t_0_ and culture fresh weights and target proteins were sampled at this time and 2, 4, 6, 8, 16, 24, 32, 48, 72 and 96 hours later. In the PTI elicited experiment two situation were differentiated: the early phase when the shift from homeostasis to what we considered to be fully induced PTI at 16 hours took place and which we termed condition 2 (C2) and the later phase when the plants were in a steady state of PTI, from 16 to 96 hours which we termed condition 3 (C3). K_d_ and K_s_ values for C3 incorporating the dilution factor determined from the culture fresh weights, which were lower than in the normally growing uninduced cultures (Supplementary Fig 3B) were calculated and classified for 69 proteins as above for C1 using the t_0_, 8 hours and all later time points. 5, 31 and 33 proteins were classified as slow, intermediate and fast, reciprocal behavior as in steady state growth, where the majority of proteins degraded slowly. The overlapping set with C1 was 57 proteins (Supplemental table 10). Of the 23 proteins in the target set that were components of the photosynthetic apparatus, 15 allowed calculations of K_d_ and K_s_ values in both conditions. Degradation rates increased significantly for these proteins in steady state PTI while synthesis rates were lower but not significantly (Fig 5A). This indicates that the decreased abundance of photosynthesis related proteins is also regulated post-transcriptionally primarily by alterations in protein turn-over.

**Figure 5:**
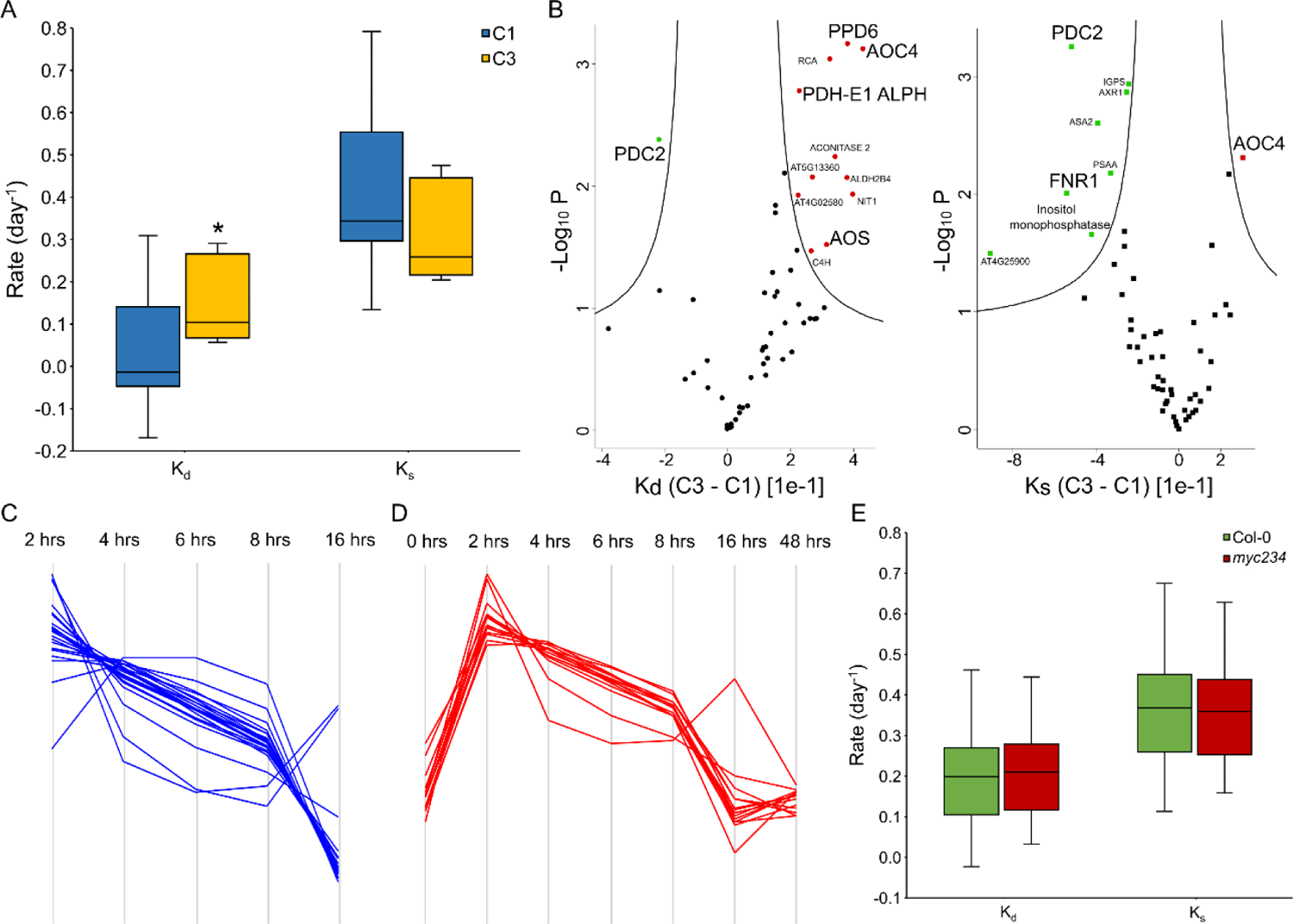
Post-transcriptional regulation of target protein abundance in flg22 induced PTI. A: Synthesis (K_s_) and degradation (K_d_) rates of photosynthesis associated proteins (PAPs) under conditions of homeostasis (C1) and fully induced PTI (C3). Star indicates statistically significant difference as determined by a two-sided student’s t-test with α=0.05 and n=15. B: Statistically significant differences in K_d_ and K_s_ values of all target proteins between conditions of fully induced PTI and homeostasis (Permutation based FDR student’s t-test with α=0.05, S0=0.1 and n=3). C: Chronological profile of protein K_s_ values in the establisher phase of PTI (C2) (n=23). D: Chronological profile of protein K_s_ values that were determined for all three conditions, homeostasis, establishment of PTI up to 16 hrs post exposure to flg22 and fully induced PTI (C1, C2 and C3) (n=14). E: K_s_ and K_d_ values of all targets in the Col-0 and myc234 backgrounds in fully induced PTI (n=53).

Several proteins all localized in the chloroplast and involved in photosynthesis and or metabolism showed significant changes in turn-over rates (K_d_ and/or K_s_ values) upon flg22 exposure (Fig 5B). The photosystem II oxygen evolving complex PsbP family protein PPD6 and the pyruvate dehydrogenase E1-alpha subunit (PDH-E1-Alpha) were both degraded more rapidly. Pyruvate decarboxylase 2 (PDC2) which has been shown to render plants more tolerant to hypoxia (*59*) was turned-over more slowly in total (smaller K_d_ and K_s_ values). Inositol monophosphatase (AT5G64380) was synthesized more slowly. FNR1, the terminal protein in the electron transfer chain, whose decreased abundance in induction of PTI is not known to be regulated by way of changes in protein turn-over also experienced a decrease in its synthesis rate in response to flg22.

Also, the enzymes catalyzing the last two reactions in 12-oxophytodienoic acid (OPDA) synthesis in the JA biosynthesis pathway, allene oxide synthase 4 (AOS) and allene oxide cyclase (AOC4) had greater K_d_ values in PTI. AOC4 also showed higher synthesis rates. This phytohormone is known to be suppressed by SA in the immune response to biotrophic pathogens and the abundance of these key enzymes in its biosynthesis pathways is not known to be regulated post-transcriptionally. Other proteins with turn-over rates affected by PAMP exposure are shown in Fig 5B.

C2 K_d_ values were calculated with alternative equation 7 that incorporate the protein abundance fold changes over time. K_s_ values were calculated as for conditions C1 and C3. These were acquired for 2, 4, 6, 8 and 16 hours post flg22 exposure as related to uninduced samples by model fitting the PQI values we measured at 1, 3 and 16 hours in the PRM experiments described above. K_d_ and K_s_ values were only calculated for proteins with FCPs higher than 1.5 or lower than 0.75 amounting to a total of 52 proteins. Values, model equations and correlation (R^2^) measures are given in Supplementary table 11. The mean correlation was 0.95 indicating high quality of fit for all proteins. After further quality control filtering 28 proteins remained with calculated K_d_ values and K_s_ values at 2, 4, 6, 8 and 16 hours.

Hierarchical cluster analysis of these 28 proteins (Table 1) showed that the great majority (23) of them had K_s_ values that changed over time and that were at a maximum 2 hours after addition of flg22 to the growth medium, decreasing over time and returning to their lowest levels 16 hours after treatment with the PAMP (Fig 5C). Moreover, K_s_ values of 14 of the 28 were calculated in all three conditions (C1, C2 and C3) and showed a strong increase after 0 hrs (C1), declining throughout the PTI establisher phase (C2) and returning to basal levels in fully induced PTI (C3) (Fig 5D). So, K_s_ values and mRNA levels both showed similar behavior, being elevated at earlier time points at the outset of proteome remodeling, indicating both played roles in regulating protein synthesis. Synthesis rates are determined by many parameters including mRNA levels, however the correlations (R^2^) between K_s_ values and mRNA levels for 10 proteins for which mRNA levels were available were 0.03, 0.01 and 0.04 at 1, 3 and 16 hrs post PAMP exposure, respectively. This shows that in our experiment the two were largely independent and that other factors primarily determined the synthesis rate itself.

**Table 1.**
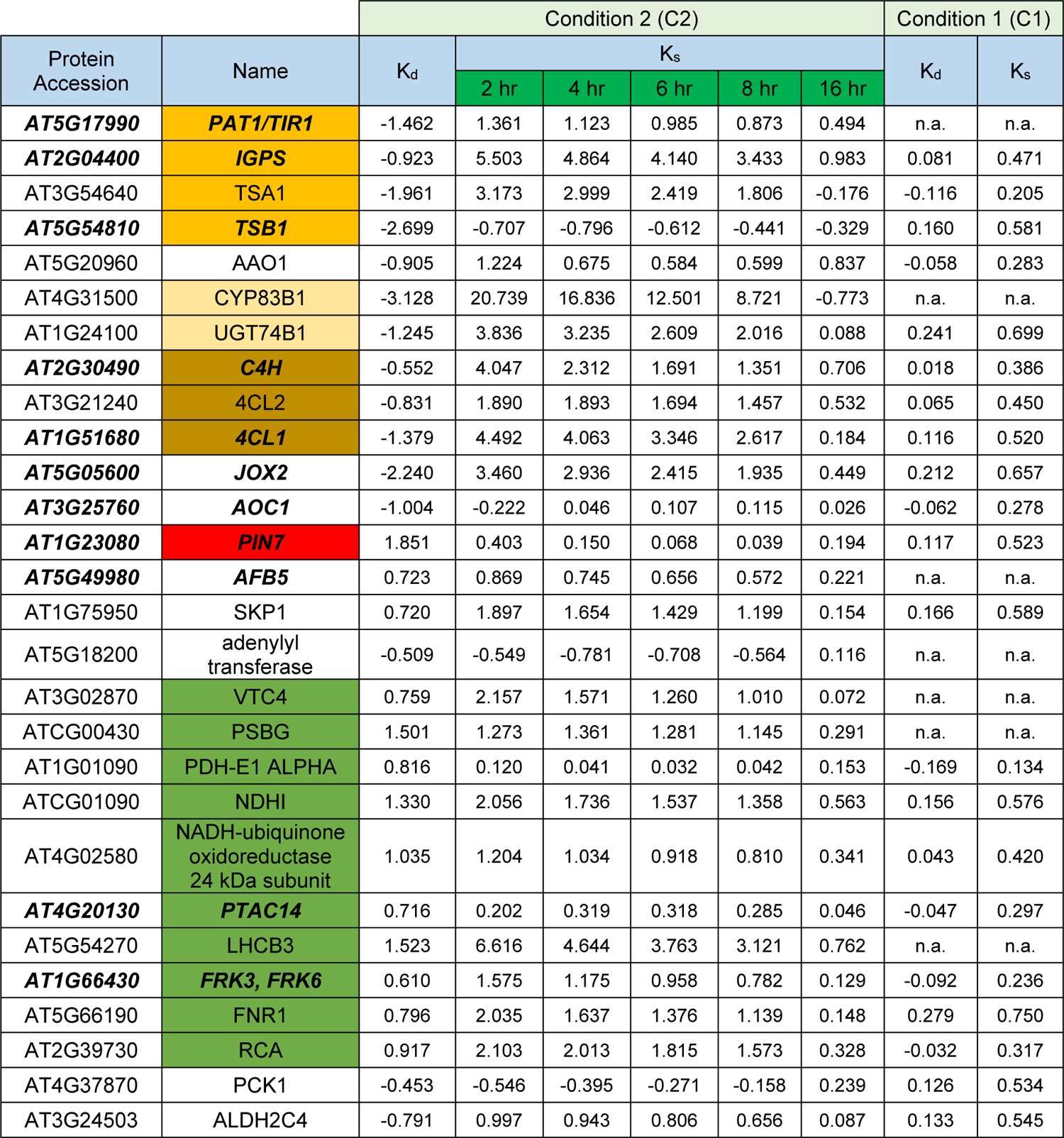
Protein synthesis (K_s_) and degradation (K_d_) rates under optimal growth conditions (C1) and the shift between growth and immunity (C2). Proteins in bold italics were also reported to be translationally regulated by (*44*)

**Table 2.**
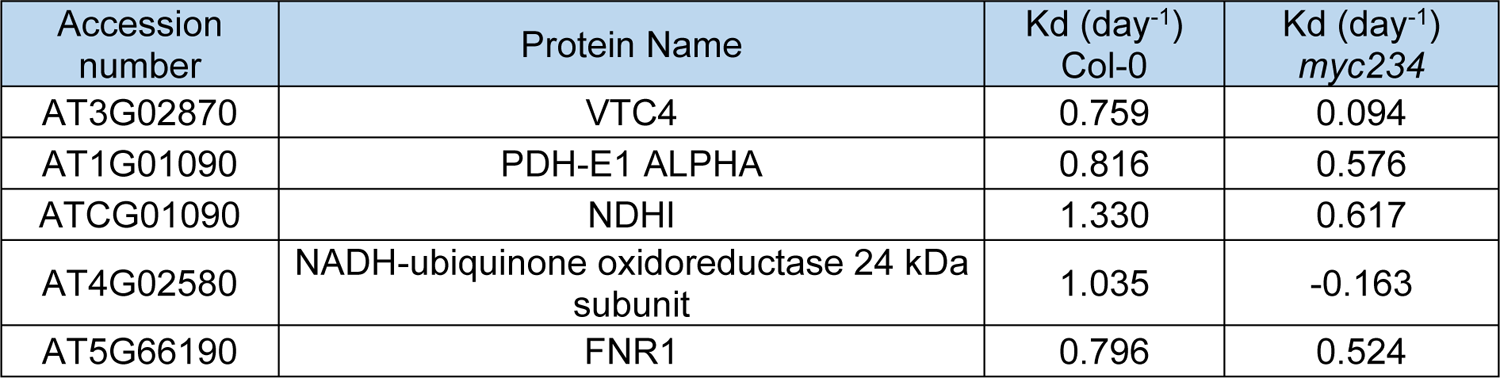
Protein synthesis and degradation rates of PAPs that were measured in both Col-0 and *myc234* backgrounds in the establisher phase (C2) up to 16 hours post flg22 perception

| Accession number | Protein Name | Kd (day<sup>-1</sup>) Col-0 | Kd (day<sup>-1</sup>) *myc234* |
| --- | --- | --- | --- |
| AT3G02870 | VTC4 | 0.759 | 0.094 |
| AT1G01090 | PDH-E1 ALPHA | 0.816 | 0.576 |
| ATCG01090 | NDHI | 1.330 | 0.617 |
| AT4G02580 | NADH-ubiquinone oxidoreductase 24 kDa subunit | 1.035 | -0.163 |
| AT5G66190 | FNR1 | 0.796 | 0.524 |

Of the 28 proteins 9 belonged to tryptophan, GS and phenylpropanoid biosynthesis pathways (Table 1 orange colors). K_d_ values of these proteins decreased even becoming negative indicating the protein pool was stabilized while at the same time K_s_ values increased markedly indicative of synthesis of new protein, concurrent with the increase in the abundance of these proteins measured in the PRM experiments. K_d_ of 10 proteins whose abundance decreased over time related to photosynthesis (Table 1 green color) increased, reflecting their degradation. This was also true for FNR1. Interestingly, K_s_ values of these proteins also increased despite the net decrease in their abundance over time showing that both degradation and synthesis i.e., turn-over regulate their abundance. The PIN7 K_d_ value increased sharply, and K_s_ values were below the optimal growth K_s_ value meaning that the rapid decrease in the abundance of this auxin transporter measured in the PRM experiments was mainly affected by post-transcriptional processes.

Our multi-omics measurements showed that the chronological changes in the abundance of these proteins playing functional roles in the shifts between growth and immunity underlie post transcriptional regulation in addition to regulation by way of changes in transcript abundance. The regulatory relationships are multi layered and we observed substantial lag times between transcript and protein synthesis. The same set of measurements was performed in the *myc234* knockout background. There were no significant differences in protein K_d_ or K_s_ values after long term induction of PTI (C3) 16 hours post elicitation with flg22 (Supplemental table 12, Fig 5E). However, K_d_ values of PAPs which could be calculated in the short-term PTI establisher phase C2 in both Col-0 and *myc234* backgrounds, disregarding their fold changes, are shown in Table 2. Degradation of these proteins, including FNR1, was substantially smaller in the mutant. This shows that the observed depletion of PAP proteins upon flg22 perception in the induction of PTI and post-transcriptional regulation thereof is dependent on MYC2 and homologs.

## Discussion

Ninety nine targets were chosen based on their amenability to MS detection, their constituency of complete biosynthesis pathways and our interest in the still understudied role of JA and auxin in PTI. Sixteen hours of PAMP treatment previously elicited a strong PTI response which we considered fully induced because of the substantial changes to the entire proteome that were measured (*21*). Transcriptomics studies employing flg22 (*20, 60*) or Met-JA treatment (*34, 61*) used varying exposure times from 1 to 16 hours and (*60*) showed that flg22 marker gene expression generally returned to basal levels between 12 and 24 hours, which we also found. The sixteen hours recovery phase in flg22 free medium was an arbitrary choice because to our knowledge no studies of this kind had been performed previously.

Both basal and induced immunity, phytohormone biosynthesis and a plethora of other metabolic and physiological processes are attuned to a 24 hour time cycle by the circadian clock, a central, essentially free running, genetic oscillator (*62*). The morning clock genes CIRCADIAN CLOCK ASSOCIATED 1 (CCA1) and LATE ELONGATED HYPOCOTYL (LHY) gate stomatal induced defenses and the expression of numerous genes pertinent to PTI shows a circadian rhythm dependent on CCA1 and the time of day of flg22 exposure (*63–65*). Our experiment uncoupled the two (clock and immunity) as evidenced by a lack of change in the protein abundance of our targets. Concerns that physical manipulation of the seedlings during the transfer between media may have elicited a JA response where similarly alleviated.

Transcripts and proteins comprising the tryptophan and aliphatic and indolic glucosinolate (IG) biosynthesis pathways increased in their abundance after exposure to flg22. The latter are important defense compounds required for callose deposition in PTI, and the elevated expression of IG biosynthesis genes has been described previously in response to flg22 (*23*). Tryptophan is the metabolic precursor channeling into IG biosynthesis and we link induction of its synthesis to flg22 perception here and in our previous study (*21*). We did not measure IG levels themselves because (*23*) showed that cleavage by the myrosinases PEN2 and PEN3 to activate them hampers direct measurement.

Induction of both tryptophan and GS biosynthesis pathways has been reported to be an integral part of JA mediated defenses in response to wounding, necrotrophic pathogens, herbivores, exogenous application of JA and constitutive JA signaling in a *jazD* mutant devoid of ten of the family members of the JAZ repressor proteins (*57, 66–70*). Introgression of a *myc234* loss of function mutation into the *jazD* background abolished gene expression of biosynthetic pathways and accumulation of tryptophan and IGs (*69*) showing dependence on these transcription factors. In addition, the *myc234* mutant is known to be compromised in both tryptophan and GS biosynthesis (*57*). This agrees with our measurements of basal levels of these pathway proteins in the triple mutant which were below wild type levels. Interestingly however, protein levels increased in the mutant as wild type upon flg22 treatment in our experiment although at 3 hours as opposed to 1 hour after exposure, suggesting that in PTI, tryptophan and GS biosynthesis seems not to depend on MYC2 and homologs. Conversely, MYC2 has been documented as a suppressor of tryptophan and IG biosynthesis gene expression following treatment of plants with exogenous Me-JA (*71*).

Both MYC (*57*) and MYB (*72, 73*) transcription factors regulate expression of GS biosynthesis genes and MYB51 downstream of ethylene signaling mediates the elevated expression of IG biosynthesis genes in PTI (*23*). So our measurements further underscore the complex often context dependent activity of signaling through MYC2 and that two genetically independent pathways control GS biosynthesis and may fine tune immunity to different pathogens and pests. One may speculate that protein abundance elevated in the triple mutant above wild type levels may be due to a lack of MYC-MYB protein interactions (*57*) which may act as a molecular switch sequestering members of one or the other transcription factor family from inducing expression of IG target genes dependent on the prevalent defense scenario.

Tryptophan is also a metabolic precursor of auxin/IAA and overexpression of TRYPTOPHAN SYNTHASE BETA subunit (TSB) genes from broccoli in *Arabidopsis* led to accumulation of both auxin and GS (*74*). Indole-3-acetaldoxime (IAOx) is the metabolic precursor of both (*75*) as well as camalexin (*76*) in *Arabidopsis* and this compound and metabolic flux have been discussed as possible modulators of the growth defense trade-off. However, this opinion has not solidified and is not supported by our measurements. Despite the induced expression of tryptophan, IG and camalexin biosynthesis pathway transcripts and proteins and a general increase in tryptophan itself (*21*) upon flg22 treatment, auxin levels were affected only slightly decreasing by about 20% throughout the experiment. Thus, a major decrease in total auxin levels does not seem to accompany PTI to affect the reduction in seedling growth which we measured upon PAMP exposure. Conspicuously, a reduction in total free auxin/IAA content was not observed in the *myc234* mutant background. *myc2* single loss of function mutants *jin1-9* and *jin1-10* also exhibited elevated free auxin levels (*77*) supporting our results that MYC2 negatively regulates these. The authors attributed this suppressive effect to the inhibition of tryptophan biosynthesis and so to reduced availability of IAA metabolic precursors.

Most tryptophan, GS and camalexin biosynthesis pathway transcripts returned to basal levels at the end of the recovery phase of the experiment, yet cognate protein levels remained elevated even 16 hours after shift of the plants to flg22 free medium. It is known that proteome remodeling is energetically costly, and that the proteome is more stable and conserved than the transcriptome (*78*) and adjustment of the abundance of these proteins may not immediately be necessary for a physiological shift back to homeostasis. Also, the perpetuated elevation may be attributed to a type of priming phenotype where the plant is prepared to deal with future attackers upon initial activation of PTI (*79*).

The abundance of the polar auxin transport proteins PIN3 and PIN7 decreased early and continuously over the period of flg22 exposure yet increased again rapidly returning to basal levels when the PAMP was removed. A decrease in the abundance of these proteins and perturbed auxin distribution has been reported in response to *Alternaria brassica* infection (*67*). The chronological profile of the abundance of these proteins measured here (decrease under and especially rapid return to normal levels in the absence of PAMP stimulus) suggests that local auxin gradients may be more important in regulating growth defense shifts than global auxin levels *per se*. PIN3 and PIN7 may be quintessential in modulating this growth defense trade-off. Their expression is not downstream of JA signaling in response to flg22 as in the immunity to necrotrophic pathogens or xylem development (*67*) because protein abundance levels in the *myc234* mutant were similar to wild type.

Down-regulation of the expression of photosynthesis associated genes (PAG) and photosynthetic activity under numerous biotic and abiotic stress as well as exogenous application of phytohormones has been amply documented ((*58, 80*) and citations therein). A reduction in the abundance of cognate proteins has been shown here and by us and others previously (*21, 40*). Two recent reports disclosed these phenomena and concurrent production of chloroplast reactive oxygen species (cROS) as part of the active plant defense response to a wide range of bacterial effectors (*81, 82*). Two types of cROS are generated, singlet oxygen (^1^O_2_) by transfer of excitation energy from triplet-state Chl in PSII (*83*) and hydrogen peroxide (H_2_O_2_) by electron transfer and oxygen reduction (Mehler reaction) from ferredoxin in PSI (*84*). Depletion of photosynthesis associated proteins (PAP) particularly PSI and PSII components and inhibition of photosynthesis were a prerequisite for cROS production, suppression of PTI and robust induction of ETI in response to pathogen effectors (*81, 82*). Blocking of electron transfer from PSII by 3-(3,4-dichlorophenyl)-1,1-dimethylurea (DCMU) led to inhibition of H_2_O_2_ production at PSI and compromised immunity against *P. syringae* DC3000 *hrpA*, a disarmed pathogen unable to deliver effectors into the cell and elicit ETI (*81*).

We have measured the reduced abundance of PAPs in response to the PAMP flg22 repeatedly although this was excluded by SU and co-workers. Presumably this was because the moderate fold changes we observed escaped detection by the SDS-PAGE technique that they used to visually show down regulation of PSI and PSII proteins. We also link the diminished abundance of PAPs in flg22 elicited PTI to MYC2 activity because it was severely diminished in the *myc234* mutant background. Attaran and co-workers (*58*) reported depletion of PAPs in response to exogenous application of Met-JA but only transient inhibition of photosynthetic activity resulting from a lack of CO_2_. This agrees with upheld electron transfer to PSI. Current models (*55*) postulate electron transfer along the thylakoid membrane in PTI leading to H_2_O_2_ production at PSI whereas extensive depletion of PAPs leads to its interruption and more vigorous ^1^O_2_ production in ETI. Both cROS are bioactive defense compounds.

FNR1 is the final protein in the electron transfer chain that has the reducing potential to transfer an electron from ferredoxin onto NADP^+^ producing NADPH. If this reaction does not proceed because of a lack of FNR1 the electron instead goes into oxygen photoreduction and H_2_O_2_ production (*81*). The *fnr1 x fnr2* heterozygote displayed reduced photosynthetic activity and increased ion leakage indicative of increased ROS production (*85*). Here we have shown a decrease in FNR1 transcript and protein abundance upon flg22 exposure. Furthermore, we have measured a significant reduction in the protein’s synthesis rate in PTI elicited by the PAMP (C3) and an elevation in its degradation rate in the earlier establisher phase of PTI (C2), which was dependent on MYC2 and homologs. Our results suggest reshaping of the photosynthetic protein apparatus as an active defense response mediated by MYC2 and homologs in PTI as opposed to the MPK3/MPK6 mediated mechanism disclosed for ETI (*82*). The molecular switch between growth and immunity itself is FNR1 which we also show is post-transcriptionally regulated by way of alterations in its synthesis and degradation rates. Thus FNR1 and indeed the entirety of PAPs may be considered potential players in the trade-off between growth and defense.

Our experiments in the *myc234* mutant uncovered a role of these bHLH transcription factors in the early establisher phase of PTI and potentially in defense against biotrophic pathogens. The abundance of target proteins did not increase 1 hour after exposure to the PAMP in this genotype as it did in the wild type. Collectively, also considering the attenuated PAP and auxin depletion in the *myc234* mutant our results suggest a positive effect of MYC2 and homologs in flg22 induced PTI.

An inhibitory effect of the transcription factors on defense against biotrophic pathogens is amply documented by way of JA antagonism of the SA pathway and indeed the triple knockout mutant hyper-accumulated SA as has also been shown previously for *jin1* (*86*). MYC2 activates transcription of NAC transcription factors in response to the JA-Ile mimic coronatine (*87, 88*) that repress transcription of *ICS1* and SA accumulation. The *jin1* mutant lacking MYC2 has been shown to be more resistant to both biotrophic and necrotrophic pathogens (*86*). On the other hand, MYC2 has also been reported to bind to the second intron of *ICS1* and induce its expression and plants over-expressing it are hyper resistant to multiple strains of virulent and avirulent bacterial pathogens (*32*). Also in the absence of PHYTOALLEXIN DEFFICIENT 4 (PAD4) MYC2 was able to induce ENHANCED DISEASE SUSCEPTIBILTY 5 (EDS5) and SA accumulation (*28*).

MYC2 and homologs are canonical JA responsive transcription factors and some of the effects we observed were also observed downstream of JA signaling. However, it may be noted that throughout our entire experiment JA and JA-Ile were hardly elevated above basal levels (low ng/g FW range) despite significant increases in fold change. These phytohormones increase around a hundred-fold of what we measured in *Arabidopsis* response to wounding (*89*) and insect herbivory (*90*). This raises the question if MYC2, MYC3 and MYC4 may be targets of another signaling pathway in *Arabidopsis* flg22 induced PTI. Liu and co-workers disclosed degradation of JAZ repressor proteins and MYC2 activation by the SA receptors NPR3 and NPR4 upon *Pseudomonas syringae* pv *maculicola* ES4326 with *avrRpt2* but not with the same strain devoid of the effector (*91*). This was evidence of non-canonical MYC2 activation in the ETI context. Our results further underscore the emerging picture of a complex relationship between hormone signaling pathways that is temporally and context dependent to fine-tune defense responses which requires reevaluation of longstanding paradigms.

We observed a positive correlation (R^2^) of around 0.4 between protein and mRNA abundance under conditions of homeostasis in both the wild type and *myc234* mutant meaning around 40% of abundance of our target proteins is explained by mRNA levels. This value finds a general consensus in the literature, but correlation can vary considerably depending on organism, tissue and biological context (*92, 93*). Indeed, correlation decreased substantially even becoming negative in the early establisher phase of PTI and throughout our experiment. Protein to transcript ratios also showed considerable chronological variation, both indicating a temporal discrepancy between the transcription of PAMP induced genes and the synthesis of cognate proteins. Although this in itself is not surprising, the lag time of one to a few hours in defense signaling may be.

A study in maize uncovered divergent protein to transcript abundance ratios in the developmental zones of the same leaf blade that were highly dependent on protein function in the respective zones (*94*). Another work in *Arabidopsis* described the protein to mRNA abundance correlation becoming negative as plants shifted from cold acclimation to de-acclimation (*95*) which the authors attributed to storage of nascent transcripts followed by subsequent translation on demand. Thus our results and others indicate post-transcriptional regulatory processes also determine protein abundance at any given time point in the cell.

Translational regulation is often essayed using Ribo-seq, a technique that precipitates genetically tagged ribosomes in the process of translation to quantify actively translated mRNA. Extensive changes in the amount of translated mRNA was found in photomorphogenic *Arabidopsis* (*96*) but translational control of plant immunity is not widely studied. Xu and co-workers (*42*) first observed increased translation of 523 transcripts upon elicitation of PTI by the PAMP elf18 determined by enrichment of purine nucleotides in 5’leader sequences. Another study reported translational upregulation of 2000 transcripts following exposure to flg22 for 1 hour (*44*) and in the ETI response to RPM1 activation, more than 6000 transcripts were detected exhibiting regulation of translation (*43*). These studies were all in *Arabidopsis* and nothing is reported in other plant species.

Differential protein abundance is a question of protein turn-over, so it underlies both synthesis and degradation. Measurements of these constants add an additional dimension to simply measuring relative changes in translated mRNA, which anyway only serves as a proxy for the protein synthesis rate ignoring degradation. Our study provides K_s_ and K_d_ values for our target proteins putting a value on these processes and shows that they change in response to flg22. This was the case for a set of proteins in fully established PTI (C3) but was even more pronounced in the transitionary phase between optimal growth (C1) and PTI (C2) shown in Table 1. Many of these proteins showed changes in both K_s_ and K_d_ values, evidence for a more elaborate relationship between the two processes in regulating their abundance over time. Eleven of the 28 proteins in Table 1 were also reported to be translationally regulated in response to flg22 by Tabassum and co-workers (*21*) whereas only 3 also responded to elf18 (*42*). Protein synthesis rates changed substantially in the earlier phases of PTI establishment often reaching a maximum 2 hour post elicitation with the PAMP. A study in human cell lines (*97*) also noted that changes in protein synthesis rates are primary determinants of protein abundance in transitions between physiological steady states. The correlation between K_s_ values and mRNA levels, where available was negligible. While our study is only a snapshot of the entire proteome it produced evidence of combined translational and post-translational regulation of protein abundance in the establishment and perpetuation of PTI and potentially defense against biotrophic pathogens. Even less is understood about the role of phytohormones in these processes. Our study gives a first perspective that MYC2 and homologs have some effect on protein synthesis and degradation of some of our targets, particularly PAPs in PTI.

## Supporting information

List of Primers Used

## Data availability

All raw and metadata have been deposited to ProteomeXchange Consortium via the Pride partner repository with the dataset identifier PXD041215 and can be found at: http://www.proteomexchange.org.

## Acknowledgments

We gratefully acknowledge the Leibniz Association for funding. Mohammad Abukhalaf was funded by DFG project grant HO 5063/2-1. We express our heartfelt gratitude to Prof. Dr. Tina Romeis for her continued support throughout the last years. We declare no conflict of interest.

## Author contributions

MAK designed experiments, performed experiments, analyzed data, disseminated data by way of data repositories and wrote the manuscript. CP and DT performed experiments. JZ performed experiments and analyzed data. WH conceived the study, acquired funding, supervised the study, designed experiments, analyzed data and wrote the manuscript.

---

**Supplementary Figure 1:**
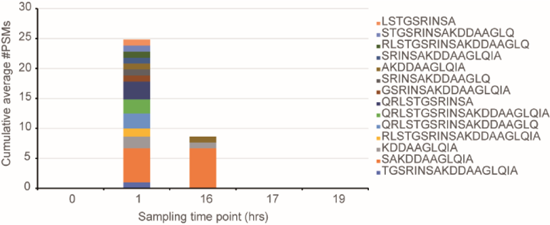
Flg22 full length and degradation products quantified at the 1, 16, 17 and 19 hrs sampling points (17 and 19 hrs represent 1 and 3 hrs post switch to flg22 free medium). Mean number of peptide spectral matches (#PSMs) (number of MS^2^ spectra annotated with a particular sequence) n=3 was used as PQI.

**Supplementary Figure 2:**
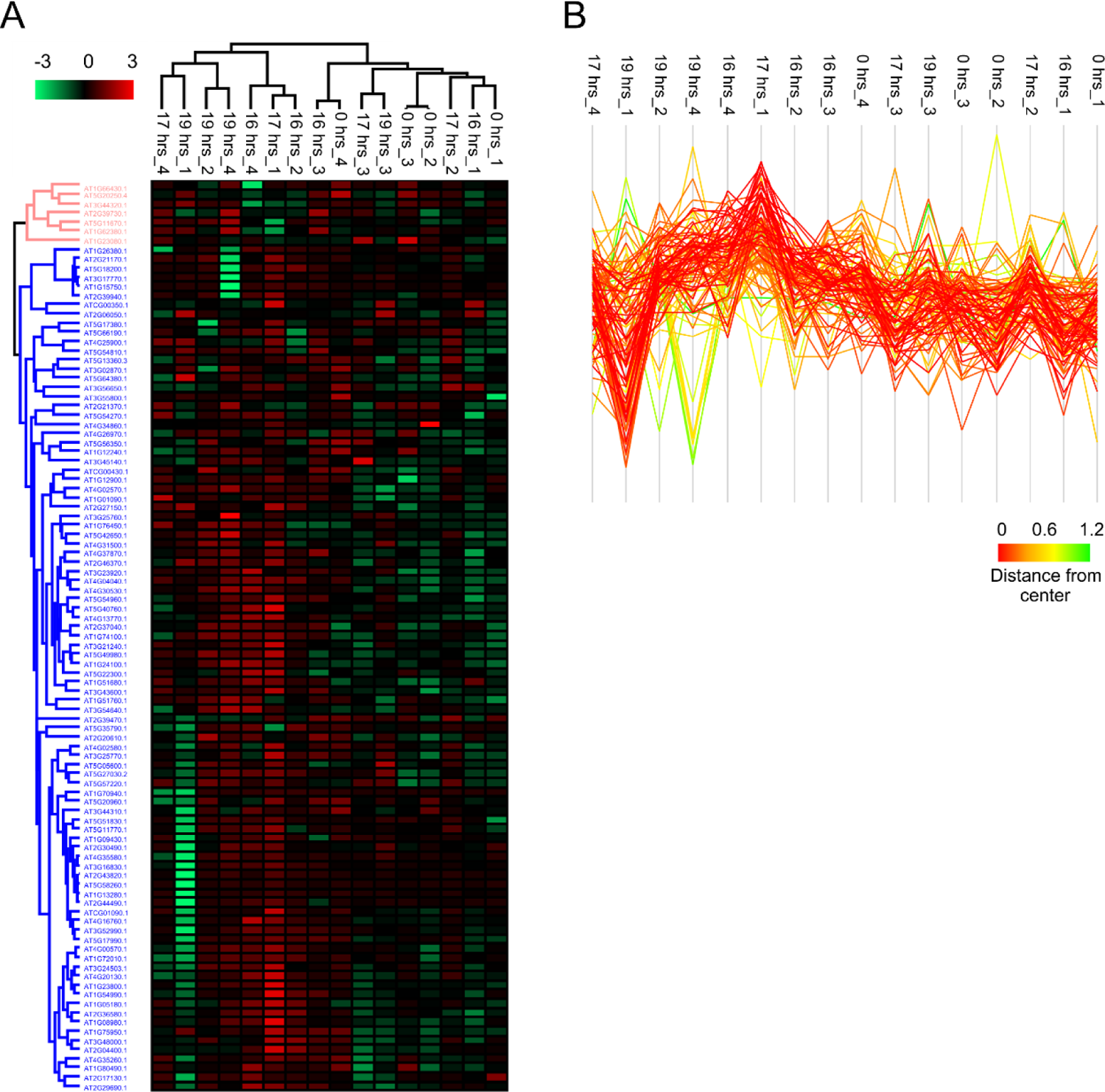
PRM based quantification of target proteins at 0, 16, 17 and 19 hrs sampling timepoints however without addition of flg22 to the medium at 0 hrs. Plants were transferred to new medium at the 16 hrs time point. A. Hierarchical cluster analysis of z-score transformed PQI values. B. Profile of larger blue cluster in A (n=92).

**Supplementary Figure 3:**
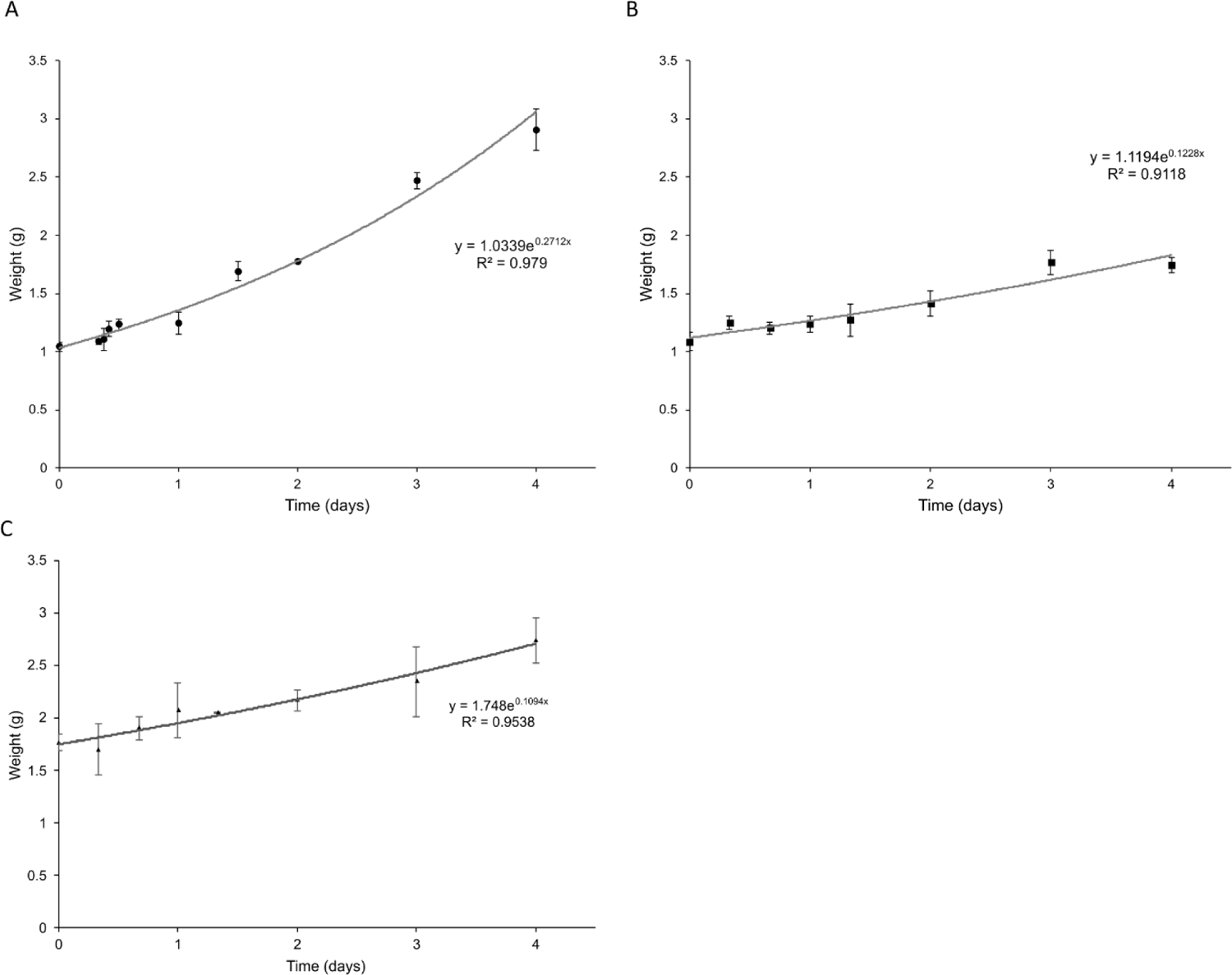
Seedling growth in culture. A. Col-0 optimal growth conditions (homeostasis). B. Col-0 flg22 replete medium (1 µM in medium). C. *myc234* flg22 replete medium.

## Notes

http://www.proteomexchange.org

### Competing Interest Statement

The authors have declared no competing interest.

## References

1. S. H. Spoel et al., NPR1 modulates cross-talk between salicylate- and jasmonate-dependent defense pathways through a novel function in the cytosol. Plant Cell 15, 760–770 (2003).

2. S. H. Spoel et al., Proteasome-mediated turnover of the transcription coactivator NPR1 plays dual roles in regulating plant immunity. Cell 137, 860–872 (2009).

3. N. Aerts, M. Pereira Mendes, S. C. M. Van Wees, Multiple levels of crosstalk in hormone networks regulating plant defense. Plant J 105, 489–504 (2021).

4. C. M. Pieterse, D. Van der Does, C. Zamioudis, A. Leon-Reyes, S. C. Van Wees, Hormonal modulation of plant immunity. Annu Rev Cell Dev Biol 28, 489–521 (2012).

5. J. Chaiwanon, W. Wang, J. Y. Zhu, E. Oh, Z. Y. Wang, Information Integration and Communication in Plant Growth Regulation. Cell 164, 1257–1268 (2016).

6. G. L. B. Gomes, K. C. Scortecci, Auxin and its role in plant development: structure, signalling, regulation and response mechanisms. Plant Biol (Stuttg) 23, 894–904 (2021).

7. X. Cao et al., The Roles of Auxin Biosynthesis YUCCA Gene Family in Plants. Int J Mol Sci 20, (2019).

8. J. Petrásek, J. Friml, Auxin transport routes in plant development. Development 136, 2675–2688 (2009).

9. I. Blilou et al., The PIN auxin efflux facilitator network controls growth and patterning in Arabidopsis roots. Nature 433, 39–44 (2005).

10. J. M. Zhou, Y. Zhang, Plant Immunity: Danger Perception and Signaling. Cell 181, 978–989 (2020).

11. L. Gómez-Gómez, T. Boller, FLS2: an LRR receptor-like kinase involved in the perception of the bacterial elicitor flagellin in Arabidopsis. Mol Cell 5, 1003–1011 (2000).

12. D. Tang, G. Wang, J. M. Zhou, Receptor Kinases in Plant-Pathogen Interactions: More Than Pattern Recognition. Plant Cell 29, 618–637 (2017).

13. E. Smakowska-Luzan et al., An extracellular network of Arabidopsis leucine-rich repeat receptor kinases. Nature 553, 342–346 (2018).

14. M. Stegmann et al., The receptor kinase FER is a RALF-regulated scaffold controlling plant immune signaling. Science 355, 287–289 (2017).

15. W. Tian et al., A calmodulin-gated calcium channel links pathogen patterns to plant immunity. Nature 572, 131–135 (2019).

16. U. Dubiella et al., Calcium-dependent protein kinase/NADPH oxidase activation circuit is required for rapid defense signal propagation. Proc Natl Acad Sci U S A 110, 8744–8749 (2013).

17. G. Bi et al., Receptor-Like Cytoplasmic Kinases Directly Link Diverse Pattern Recognition Receptors to the Activation of Mitogen-Activated Protein Kinase Cascades in Arabidopsis. Plant Cell 30, 1543–1561 (2018).

18. W. Hoehenwarter et al., Identification of novel in vivo MAP kinase substrates in Arabidopsis thaliana through use of tandem metal oxide affinity chromatography. Mol Cell Proteomics 12, 369–380 (2013).

19. T. Guerra et al., Calcium-dependent protein kinase 5 links calcium signaling with N-hydroxy-l-pipecolic acid- and SARD1-dependent immune memory in systemic acquired resistance. New Phytol 225, 310–325 (2020).

20. L. Navarro et al., The transcriptional innate immune response to flg22. Interplay and overlap with Avr gene-dependent defense responses and bacterial pathogenesis. Plant Physiol 135, 1113–1128 (2004).

21. M. Bassal et al., Reshaping of the Arabidopsis thaliana Proteome Landscape and Co-regulation of Proteins in Development and Immunity. Mol Plant 13, 1709–1732 (2020).

22. K. Maleck et al., The transcriptome of Arabidopsis thaliana during systemic acquired resistance. Nat Genet 26, 403–410 (2000).

23. N. K. Clay, A. M. Adio, C. Denoux, G. Jander, F. M. Ausubel, Glucosinolate metabolites required for an Arabidopsis innate immune response. Science 323, 95–101 (2009).

24. B. A. Halkier, J. Gershenzon, Biology and biochemistry of glucosinolates. Annu Rev Plant Biol 57, 303–333 (2006).

25. M. Burow, B. A. Halkier, How does a plant orchestrate defense in time and space? Using glucosinolates in Arabidopsis as case study. Curr Opin Plant Biol 38, 142–147 (2017).

26. Y. Peng, J. Yang, X. Li, Y. Zhang, Salicylic Acid: Biosynthesis and Signaling. Annu Rev Plant Biol 72, 761–791 (2021).

27. R. A. Hillmer et al., The highly buffered Arabidopsis immune signaling network conceals the functions of its components. PLoS Genet 13, e1006639 (2017).

28. A. Mine et al., An incoherent feed-forward loop mediates robustness and tunability in a plant immune network. EMBO Rep 18, 464–476 (2017).

29. K. Tsuda, M. Sato, T. Stoddard, J. Glazebrook, F. Katagiri, Network properties of robust immunity in plants. PLoS Genet 5, e1000772 (2009).

30. N. Hatsugai et al., A plant effector-triggered immunity signaling sector is inhibited by pattern-triggered immunity. EMBO J 36, 2758–2769 (2017).

31. K. Kazan, J. M. Manners, MYC2: the master in action. Mol Plant 6, 686–703 (2013).

32. J. K. Gautam, M. K. Giri, D. Singh, S. Chattopadhyay, A. K. Nandi, MYC2 influences salicylic acid biosynthesis and defense against bacterial pathogens in Arabidopsis thaliana. Physiol Plant 173, 2248–2261 (2021).

33. M. Altmann et al., Extensive signal integration by the phytohormone protein network. Nature 583, 271–276 (2020).

34. M. Zander et al., Integrated multi-omics framework of the plant response to jasmonic acid. Nat Plants 6, 290–302 (2020).

35. F. Wu et al., Mediator Subunit MED25 Couples Alternative Splicing of. Plant Cell 32, 429–448 (2020).

36. J. E. Moreno et al., Negative feedback control of jasmonate signaling by an alternative splice variant of JAZ10. Plant Physiol 162, 1006–1017 (2013).

37. Y. Liu et al., MYC2 Regulates the Termination of Jasmonate Signaling via an Autoregulatory Negative Feedback Loop. Plant Cell 31, 106–127 (2019).

38. J. M. Chico et al., CUL3. Proc Natl Acad Sci U S A 117, 6205–6215 (2020).

39. V. Marquis et al., Stress- and pathway-specific impacts of impaired jasmonoyl-isoleucine (JA-Ile) catabolism on defense signalling and biotic stress resistance. Plant Cell Environ 43, 1558–1570 (2020).

40. V. Göhre, A. M. Jones, J. Sklenář, S. Robatzek, A. P. Weber, Molecular crosstalk between PAMP-triggered immunity and photosynthesis. Mol Plant Microbe Interact 25, 1083–1092 (2012).

41. R. Schoenheimer, S. Ratner, D. Rittenberg, THE PROCESS OF CONTINUOUS DEAMINATION AND REAMINATION OF AMINO ACIDS IN THE PROTEINS OF NORMAL ANIMALS. Science 89, 272–273 (1939).

42. G. Xu et al., Global translational reprogramming is a fundamental layer of immune regulation in plants. Nature 545, 487–490 (2017).

43. L. V. Meteignier et al., Translatome analysis of an NB-LRR immune response identifies important contributors to plant immunity in Arabidopsis. J Exp Bot 68, 2333–2344 (2017).

44. N. Tabassum et al., Phosphorylation-dependent control of an RNA granule-localized protein that fine-tunes defence gene expression at a post-transcriptional level. Plant J 101, 1023–1039 (2020).

45. Y. Zhang et al., Proteome scale turnover analysis in live animals using stable isotope metabolic labeling. Anal Chem 83, 1665–1672 (2011).

46. C. J. Nelson, L. Li, A. H. Millar, Quantitative analysis of protein turnover in plants. Proteomics 14, 579–592 (2014).

47. L. Li et al., Protein Degradation Rate in Arabidopsis thaliana Leaf Growth and Development. Plant Cell 29, 207–228 (2017).

48. C. J. Nelson, R. Alexova, R. P. Jacoby, A. H. Millar, Proteins with high turnover rate in barley leaves estimated by proteome analysis combined with in planta isotope labeling. Plant Physiol 166, 91–108 (2014).

49. D. Lyon et al., Drought and Recovery: Independently Regulated Processes Highlighting the Importance of Protein Turnover Dynamics and Translational Regulation in Medicago truncatula. Mol Cell Proteomics 15, 1921–1937 (2016).

50. M. C. Chambers et al., A cross-platform toolkit for mass spectrometry and proteomics. Nature Biotechnology 30, 918–920 (2012).

51. D. Lyon et al., Automated protein turnover calculations from 15N partial metabolic labeling LC/MS shotgun proteomics data. PLoS One 9, e94692 (2014).

52. P. Majovsky et al., Targeted proteomics analysis of protein degradation in plant signaling on an LTQ-Orbitrap mass spectrometer. J Proteome Res 13, 4246–4258 (2014).

53. T. Meindl, T. Boller, G. Felix, The bacterial elicitor flagellin activates its receptor in tomato cells according to the address-message concept. Plant Cell 12, 1783–1794 (2000).

54. M. Wilhelm et al., Mass-spectrometry-based draft of the human proteome. Nature 509, 582–587 (2014).

55. G. R. Littlejohn, S. Breen, N. Smirnoff, M. Grant, Chloroplast immunity illuminated. New Phytol 229, 3088–3107 (2021).

56. P. Fernández-Calvo et al., The Arabidopsis bHLH transcription factors MYC3 and MYC4 are targets of JAZ repressors and act additively with MYC2 in the activation of jasmonate responses. Plant Cell 23, 701–715 (2011).

57. F. Schweizer et al., Arabidopsis basic helix-loop-helix transcription factors MYC2, MYC3, and MYC4 regulate glucosinolate biosynthesis, insect performance, and feeding behavior. Plant Cell 25, 3117-3132 (2013).

58. E. Attaran et al., Temporal Dynamics of Growth and Photosynthesis Suppression in Response to Jasmonate Signaling. Plant Physiol 165, 1302–1314 (2014).

59. I. Ventura et al., Arabidopsis phenotyping reveals the importance of alcohol dehydrogenase and pyruvate decarboxylase for aerobic plant growth. Sci Rep 10, 16669 (2020).

60. C. Denoux et al., Activation of defense response pathways by OGs and Flg22 elicitors in Arabidopsis seedlings. Mol Plant 1, 423–445 (2008).

61. R. Hickman et al., Architecture and Dynamics of the Jasmonic Acid Gene Regulatory Network. Plant Cell 29, 2086–2105 (2017).

62. H. Lu, C. R. McClung, C. Zhang, Tick Tock: Circadian Regulation of Plant Innate Immunity. Annu Rev Phytopathol 55, 287–311 (2017).

63. W. Wang et al., Timing of plant immune responses by a central circadian regulator. Nature 470, 110–114 (2011).

64. V. Bhardwaj, S. Meier, L. N. Petersen, R. A. Ingle, L. C. Roden, Defence responses of Arabidopsis thaliana to infection by Pseudomonas syringae are regulated by the circadian clock. PLoS One 6, e26968 (2011).

65. C. Korneli, S. Danisman, D. Staiger, Differential control of pre-invasive and post-invasive antibacterial defense by the Arabidopsis circadian clock. Plant Cell Physiol 55, 1613–1622 (2014).

66. P. Reymond et al., A conserved transcript pattern in response to a specialist and a generalist herbivore. Plant Cell 16, 3132–3147 (2004).

67. L. Qi et al., Arabidopsis thaliana plants differentially modulate auxin biosynthesis and transport during defense responses to the necrotrophic pathogen Alternaria brassicicola. New Phytol 195, 872–882 (2012).

68. H. Frerigmann et al., Regulation of Pathogen-Triggered Tryptophan Metabolism in Arabidopsis thaliana by MYB Transcription Factors and Indole Glucosinolate Conversion Products. Mol Plant 9, 682–695 (2016).

69. Q. Guo, I. T. Major, G. Kapali, G. A. Howe, MYC transcription factors coordinate tryptophan-dependent defence responses and compromise seed yield in Arabidopsis. New Phytol 236, 132–145 (2022).

70. D. J. Kliebenstein, J. Kroymann, T. Mitchell-Olds, The glucosinolate-myrosinase system in an ecological and evolutionary context. Curr Opin Plant Biol 8, 264–271 (2005).

71. B. Dombrecht et al., MYC2 differentially modulates diverse jasmonate-dependent functions in Arabidopsis. Plant Cell 19, 2225–2245 (2007).

72. M. Y. Hirai et al., Omics-based identification of Arabidopsis Myb transcription factors regulating aliphatic glucosinolate biosynthesis. Proc Natl Acad Sci U S A 104, 6478–6483 (2007).

73. T. Gigolashvili et al., The transcription factor HIG1/MYB51 regulates indolic glucosinolate biosynthesis in Arabidopsis thaliana. Plant J 50, 886–901 (2007).

74. R. Li et al., Overexpressing broccoli tryptophan biosynthetic genes BoTSB1 and BoTSB2 promotes biosynthesis of IAA and indole glucosinolates. Physiol Plant 168, 174–187 (2020).

75. S. Sugawara et al., Biochemical analyses of indole-3-acetaldoxime-dependent auxin biosynthesis in Arabidopsis. Proc Natl Acad Sci U S A 106, 5430–5435 (2009).

76. E. Glawischnig, B. G. Hansen, C. E. Olsen, B. A. Halkier, Camalexin is synthesized from indole-3-acetaldoxime, a key branching point between primary and secondary metabolism in Arabidopsis. Proc Natl Acad Sci U S A 101, 8245–8250 (2004).

77. C. F. Huang et al., Elevated auxin biosynthesis and transport underlie high vein density in C. Proc Natl Acad Sci U S A 114, E6884–E6891 (2017).

78. J. M. Laurent et al., Protein abundances are more conserved than mRNA abundances across diverse taxa. Proteomics 10, 4209–4212 (2010).

79. M. van Hulten, M. Pelser, L. C. van Loon, C. M. Pieterse, J. Ton, Costs and benefits of priming for defense in Arabidopsis. Proc Natl Acad Sci U S A 103, 5602–5607 (2006).

80. D. D. Bilgin et al., Biotic stress globally downregulates photosynthesis genes. Plant Cell Environ 33, 1597–1613 (2010).

81. M. de Torres Zabala et al., Chloroplasts play a central role in plant defence and are targeted by pathogen effectors. Nat Plants 1, 15074 (2015).

82. J. Su et al., Active photosynthetic inhibition mediated by MPK3/MPK6 is critical to effector-triggered immunity. PLoS Biol 16, e2004122 (2018).

83. P. M. Mullineaux, M. Exposito-Rodriguez, P. P. Laissue, N. Smirnoff, ROS-dependent signalling pathways in plants and algae exposed to high light: Comparisons with other eukaryotes. Free Radic Biol Med 122, 52–64 (2018).

84. D. V. Vetoshkina, B. N. Ivanov, S. A. Khorobrykh, I. I. Proskuryakov, M. M. Borisova-Mubarakshina, Involvement of the chloroplast plastoquinone pool in the Mehler reaction. Physiol Plant 161, 45–55 (2017).

85. M. Lintala et al., Depletion of leaf-type ferredoxin-NADP(+) oxidoreductase results in the permanent induction of photoprotective mechanisms in Arabidopsis chloroplasts. Plant J 70, 809–817 (2012).

86. A. Nickstadt et al., The jasmonate-insensitive mutant jin1 shows increased resistance to biotrophic as well as necrotrophic pathogens. Mol Plant Pathol 5, 425–434 (2004).

87. X. Y. Zheng et al., Coronatine promotes Pseudomonas syringae virulence in plants by activating a signaling cascade that inhibits salicylic acid accumulation. Cell Host Microbe 11, 587–596 (2012).

88. S. Gimenez-Ibanez et al., JAZ2 controls stomata dynamics during bacterial invasion. New Phytol 213, 1378–1392 (2017).

89. G. Glauser et al., Spatial and temporal dynamics of jasmonate synthesis and accumulation in Arabidopsis in response to wounding. J Biol Chem 283, 16400–16407 (2008).

90. M. K. Meena et al., The Ca2+ Channel CNGC19 Regulates Arabidopsis Defense Against Spodoptera Herbivory. Plant Cell 31, 1539–1562 (2019).

91. L. Liu et al., Salicylic acid receptors activate jasmonic acid signalling through a non-canonical pathway to promote effector-triggered immunity. Nat Commun 7, 13099 (2016).

92. C. Vogel, E. M. Marcotte, Insights into the regulation of protein abundance from proteomic and transcriptomic analyses. Nat Rev Genet 13, 227–232 (2012).

93. C. Buccitelli, M. Selbach, mRNAs, proteins and the emerging principles of gene expression control. Nat Rev Genet 21, 630–644 (2020).

94. L. Ponnala, Y. Wang, Q. Sun, K. J. van Wijk, Correlation of mRNA and protein abundance in the developing maize leaf. Plant J 78, 424–440 (2014).

95. K. Nakaminami et al., Analysis of differential expression patterns of mRNA and protein during cold-acclimation and de-acclimation in Arabidopsis. Mol Cell Proteomics 13, 3602–3611 (2014).

96. M. J. Liu et al., Translational landscape of photomorphogenic Arabidopsis. Plant Cell 25, 3699–3710 (2013).

97. A. R. Kristensen, J. Gsponer, L. J. Foster, Protein synthesis rate is the predominant regulator of protein expression during differentiation. Mol Syst Biol 9, 689 (2013).

